# Functionalized microcarriers improve T cell manufacturing by facilitating migratory memory T cell production and increasing CD4/CD8 ratio

**DOI:** 10.1101/646760

**Authors:** Nathan J. Dwarshuis, Hannah W. Song, Anokhi Patel, Theresa Kotanchek, Krishnendu Roy

## Abstract

Adoptive cell therapies (ACT) using chimeric antigen receptor (CAR) T cells have shown promise in treating cancer, but manufacturing large numbers of high quality cells remains challenging. Critically, current T cell expansion technologies only partially recapitulate the *in vivo* microenvironment found in the human lymph nodes. In these organs, T cells expand at high cell density with autocrine/paracrine signaling, as well as signals from the extracellular matrix (ECM). Here we describe a T cell expansion system using degradable gelatin microcarriers functionalized with anti-CD3 and anti-CD28 monoclonal antibodies (mAbs), which address several of these shortcomings. We show that using this system, we can achieve approximately 2-fold greater expansion compared to functionalized magnetic beads, the current industry standard. Furthermore, carriers generated higher numbers of CCR7+CD62L+ migratory, central memory T cells and CD4+ T cells across multiple donors. Both these phenotypes have emerged as important for establishing durable and effective responses in patients receiving T cell immunotherapies. We further demonstrate that carriers can achieve greater memory cell yield compared to beads across a range of IL2 concentrations from 20 U/mL to 100 U/mL. These differences were greater at lower IL2 concentrations, indicating that the carriers are more efficient. We optimized this system using a design of experiments (DOE) approach and found that the carrier concentration affects the memory cell yield in a quadratic manner, where high or low concentrations are detrimental to memory formation. Finally, we show that carriers do not hinder CAR transduction and can maintain the CD4 and memory phenotype advantages in CAR-transduced T cells.

## Introduction

T cell-based immunotherapies have received great interest from clinicians and industry due to their potential to treat, and often finally cure, cancer and other diseases [1, 2]. In 2017, Novartis and Kite Pharma acquired FDA approval for *Kymriah* and *Yescarta* respectively, two genetically-modified CAR T cell therapies against B cell malignancies. Despite these successes, CAR T cell therapies are constrained by an expensive and difficult-to-scale manufacturing process [3, 4].

State-of-the-art manufacturing techniques focus only on anti-CD3 and anti-CD28 activation, typically presented on a microbead (Invitrogen Dynabead, Miltenyi MACS beads) or nanobead (Miltenyi TransACT), but also in soluble forms in the case of antibody tetramers (Expamer) [3, 5–7]. These strategies overlook many of the signaling components present in the secondary lymphoid organs where T cells normally expand. Typically, T cells are activated under close cell-cell contact via antigen presenting cells (APCs) such as dendritic cells (DCs), which present peptide-major histocompatibility complexs (MHCs) to T cells as well as a variety of other costimulatory signals. These close quarters allow for efficient autocrine/paracrine signaling among the expanding T cells, which secrete IL2 and other cytokines to assist their own growth. Additionally, the lymphoid tissues are comprised of ECM components such as collagen, which provide signals to upregulate proliferation, cytokine production, and pro-survival pathways [8–11].

A variety of solutions have been proposed to make the T cell expansion process more physiological. One strategy is to use modified feeder cell cultures to provide activation signals similar to those of DCs [12]. While this has the theoretical capacity to mimic many components of the lymph node, it is hard to reproduce on a large scale due to the complexity and inherent variability of using cell lines in a fully Good Manufacturing Practices (GMP)-compliant manner. Others have proposed biomaterials-based solutions to circumvent this problem, including lipid-coated microrods [13], 3D-scaffolds via either Matrigel [14] or 3d-printed lattices [15], ellipsoid beads [16], and mAb-conjugated polydimethylsiloxane (PDMS) beads [17] that respectively recapitulate the cellular membrane, large interfacial contact area, 3D-structure, or soft surfaces T cells normally experience *in vivo*. While these have been shown to provide superior expansion compared to traditional microbeads, no method has been able to show preferential expansion of functional memory and CD4 T cell populations. Generally, T cells with a lower differentiation state such as memory cells have been shown to provide superior anti-tumor potency, presumably due to their higher potential to replicate, migrate, and engraft, leading to a long-term, durable response [18–21]. Likewise, CD4 T cells are similarly important to anti-tumor potency due to their cytokine release properties and ability to resist exhaustion [22, 23]. Therefore, methods to increase memory and CD4 T cells in the final product are needed, a critical consideration being ease of translation to industry and ability to interface with scalable systems such as bioreactors.

Here we describe a method using porous microcarriers functionalized with anti-CD3 and anti-CD28 mAbs for use in T cell expansion cultures. Microcarriers have historically been used throughout the bioprocess industry for adherent cultures such as stem cells and Chinese hamster ovary (CHO) cells, but not with suspension cells such as T cells [24, 25]. The carriers used in this study have a macroporous structure that allows T cells to grow inside and along the surface, providing ample cell-cell contact for enhanced autocrine and paracrine signaling. Furthermore, the carriers are composed of gelatin, which is a collagen derivative and therefore has adhesion domains that are also present within the lymph nodes. Finally, the 3D surface of the carriers provides a larger contact area for T cells to interact with the mAbs relative to beads; this may better emulate the large contact surface area that occurs between T cells and DCs. We show that compared to traditional beads, carrier-expanded T cells not only provide superior expansion, but consistently provide a higher frequency of memory and CD4 T cells (CCR7+CD62L+) across multiple donors. We also demonstrate functional cytotoxicity using a CD19 CAR. Our results indicate that functionalized microcarriers provide a robust and scalable platform for manufacturing therapeutic T cells with higher memory phenotype and CD4+ cell content.

## Results

### Microcarriers can provide greater expansion potential compared to beads

Two types of carriers, Cultispher-S (CuS) and Cultispher-G (CuG), were covalently conjugated with varying concentration of sulfo-NHS-biotin (SNB) and then coated with streptavidin (STP) (Figs. 1a and 1b). We chose to continue with the CuS carriers, which showed higher overall STP binding compared to CuG. We further set the amount of SNB to the lowest concentration per mass of carriers (5 M/g) that achieved maximal STP binding. We further verified that the carriers had active biotin binding sites at this concentration (Fig. 1c), and demonstrated that they were evenly coated throughout their interior using FITC-biotin (Fig. 1d). Finally, we confirmed that biotinylated mAbs were bound to the carriers by staining either STP or mAb-coated carriers with anti-mouse immunoglobulin G (IgG)-FITC and imaging on a confocal microscope (Fig. 1e).

**Figure 1:**
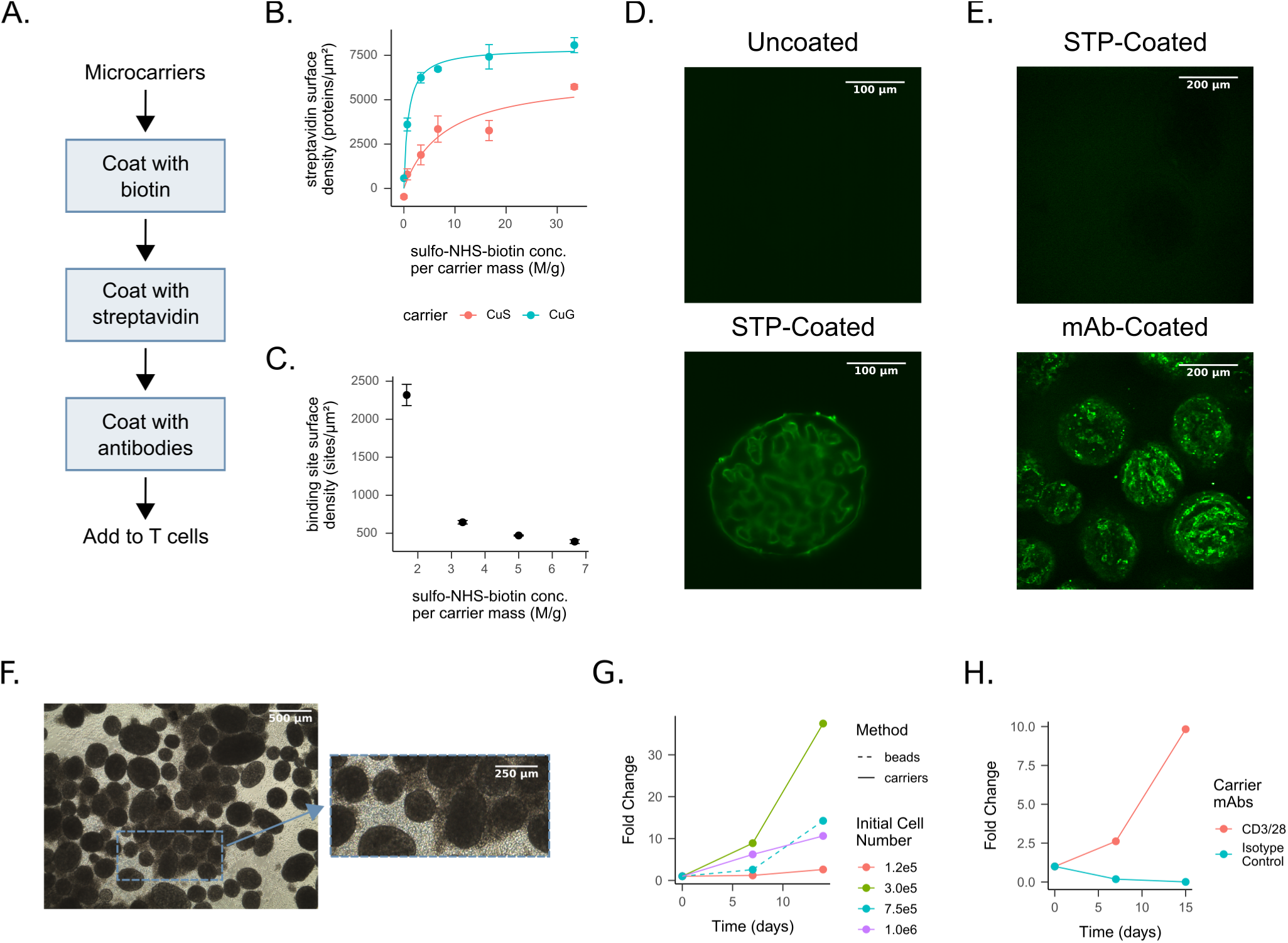
Functionalized carriers can expand T cells. a) Overview of the carrier coating process. b) Mass of bound STP vs. amount of SNB for CuS and CuG. c) Number of STP binding sites (defined using FITC-biotin) vs. amount of SNB used to biotinylate the CuS carriers. d) Lightsheet image of uncoated or STP-coated carriers stained with FITC-biotin. e) Confocal image of STP or mAb-coated carriers stained with anti-mouse IgG-FITC. f) Phase image of T cells expanded on carriers after 9 d. g) T cells expanded using either beads or carriers with three initial cell densities for 14 d. h) T cells expanded using either carriers coated with anti-CD3/anti-CD28 mAbs or isotype control. Abbreviations: streptavidin (STP); Cultispher-S (CuS); Cultispher-G (CuG); sulfo-NHS-biotin (SNB)

We next sought to determine how our functionalized carriers could expand T cells compared to state-of-the-art methods used in industry. We compared the carriers alongside traditional microbeads (Miltenyi-Biotec) by expanding T cells for 14 d with several different seeding densities in the carrier cultures (Fig. 1f). All bead expansions were performed as per the manufacturer’s protocol. We observed a higher fold change in the carrier cultures with an intermediate density of 3 *×* 10^5^ cells/mL, implying that carriers could be used to achieve greater expansion than conventional beads (Fig. 1g). We also observed no T cell expansion using carriers coated with an isotype control mAb compared to carriers coated with anti-CD3/anti-CD28 mAbs (Fig. 1h), confirming specificity of the expansion method.

We also asked how many cells were inside the carriers vs. free-floating in suspension and/or loosely attached to the surface. After seeding carriers at different densities and expanding for 14 d, we filtered the carriers out of the cell suspension and digested them using dispase to free any cells attached on the inner surface. We observed that 15 % of the total cells on day 14 were on the interior surface of the carriers (Fig. S1a), and this did not significantly change with initial seeding density (Table S1). We qualitatively verified the presence of cells inside the carriers using an 3-(4,5-dimethylthiazol-2-yl)-2,5-diphenyltetrazolium bromide (MTT) stain to opaquely mark cells and enable visualization on a brightfield microscope (Fig. S1b).

### Microcarriers produce higher frequencies of memory T cells and CD4+ T cells compared to conventional microbeads

After observing differences in expansion, we further hypothesized that the carrier cultures could lead to a different T cell phenotype. In particular, we were interested in the formation of memory T cells, as these represent a subset with higher replicative potential and therefore improved clinical prognosis [20, 21]. We measured memory T cell frequency staining for CCR7 and CD62L (both of which are present on lower differentiated T cells such as central memory cells and stem memory cells [19]). After expanding T cells for 14 days using either beads or carriers, we noted a distinctly larger frequency of memory T cells (CD62L+CCR7+) in the carrier cultures compared to the bead cultures (Fig. 2a).

**Figure 2:**
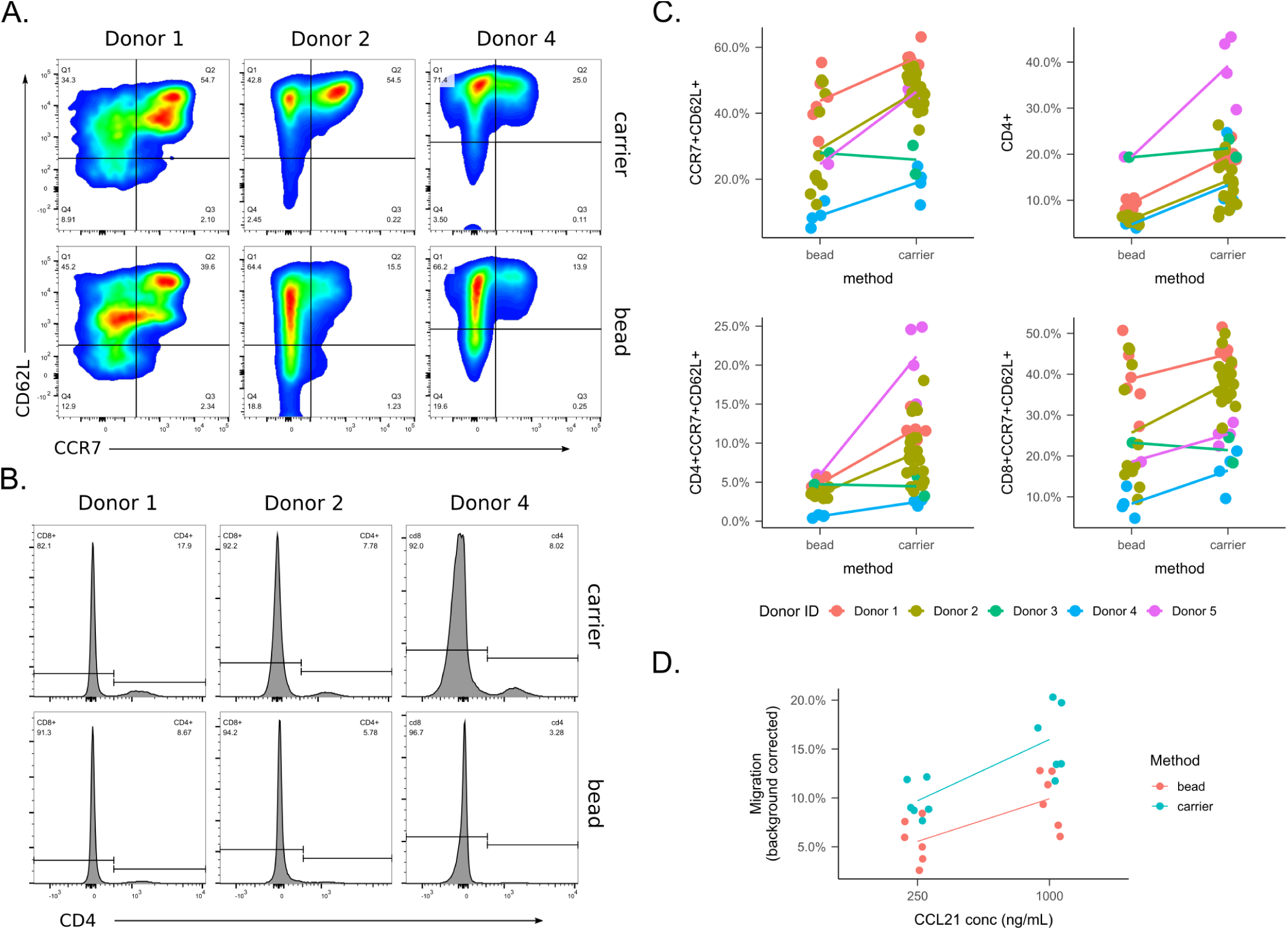
Carriers selectively expand memory T cell relative to beads across multiple donors and experiments. Note that Donors 1, 3, and 4 were CAR-transduced, and Donors 3, 4, and 5 were peripheral blood mononuclear cells (PBMCs) sorted through a magnetic activated cell sorting (MACS) column. a) Representative flow plots from three donors of memory T cells expanded for 14 d. b) Representative flow plots from three donors of CD4+ T cells expanded after 14 d. c) Fraction of memory cells obtained after 14 d for multiple donors across multiple experiments. d) T cell chemotaxis measured after 14 d expansion using a transwell with a CCL21 gradient

Of additional interest to us was the preservation of the CD4 compartment. In healthy donor samples (such as those used here), the typical CD4:CD8 ratio is 2:1. We noted that carriers produced a more balanced culture with CD4 T cells at a higher frequency compared to bead cultures, which had nearly 90 % CD8 T cells (Fig. 2b). While both systems had a preference for expanding CD8 T cells, our results indicated that the carriers allow a better CD4:CD8 ratio.

To test if this observation was consistent across experimental conditions, we pooled data for multiple T cells expansions where these markers where measured on day 14; these experiments had varying conditions as well as different donors (Table S7), and thus the pooled dataset provided a test for robustness. Comparing beads and carriers for both memory and CD4 T cell percentage, we noted that carriers provided a higher percentage in nearly every case (Fig. 2c). This trend was similar for both memory CD4+ and CD8+ subpopulations.

We analyzed this pooled dataset using linear regression analysis to determine if there was a significant difference between the beads and carriers in either memory or CD4 phenotype (Tables S2 and S3 and Figs. S2a and S2b). In both cases, the activation method (carrier vs. bead) was highly significant, with the carriers producing 13 % and 21 % greater frequencies of memory and CD4+ T cells respectively. The regression analysis also revealed that both phenotypes depended highly on donor but not on any of the aggregate process parameters (total IL2, total added glucose as calculated by the amount of media added, and the fold change of culture volume increase).

We also verified that expanded T cells were migratory using a chemotaxis assay for CCL21; since carriers produced a larger percentage of memory T cells (which have CCR7, the receptor for CCL21) we would expect higher migration in carrier-expanded cells vs. their bead counterparts. Indeed, we noted a significantly higher percentage of migration in carriers and a dose-dependent response to CCL21 (Figs. 2d and S3 and Table S6).

### Microcarriers require less IL2 for robust expansion compared to beads

We next asked if T cells required less exogenous IL2 in the carriers vs. traditional beads, as the initial hypothesis for the design was that the macroporous structure would allow more efficient autocrine and paracrine signaling by increasing local cell density. We expanded T cells using either beads or carriers for 14 d using varying amounts of IL2 from 0 U/mL to 100 U/mL added every two days. Overall, we noted that the carriers expanded the T cells more robustly (Fig. 3a) and required lower IL2 concentrations while maintaining equivalent expansion (Fig. 3b). When comparing the robustness of memory cell production, the carriers also produced more cells with less IL2 (Fig. 3c). Furthermore, the frequency of memory T cells was greater at lower IL2 concentrations in the carrier-expanded population than that of the beads (Fig. 3c).

**Figure 3:**
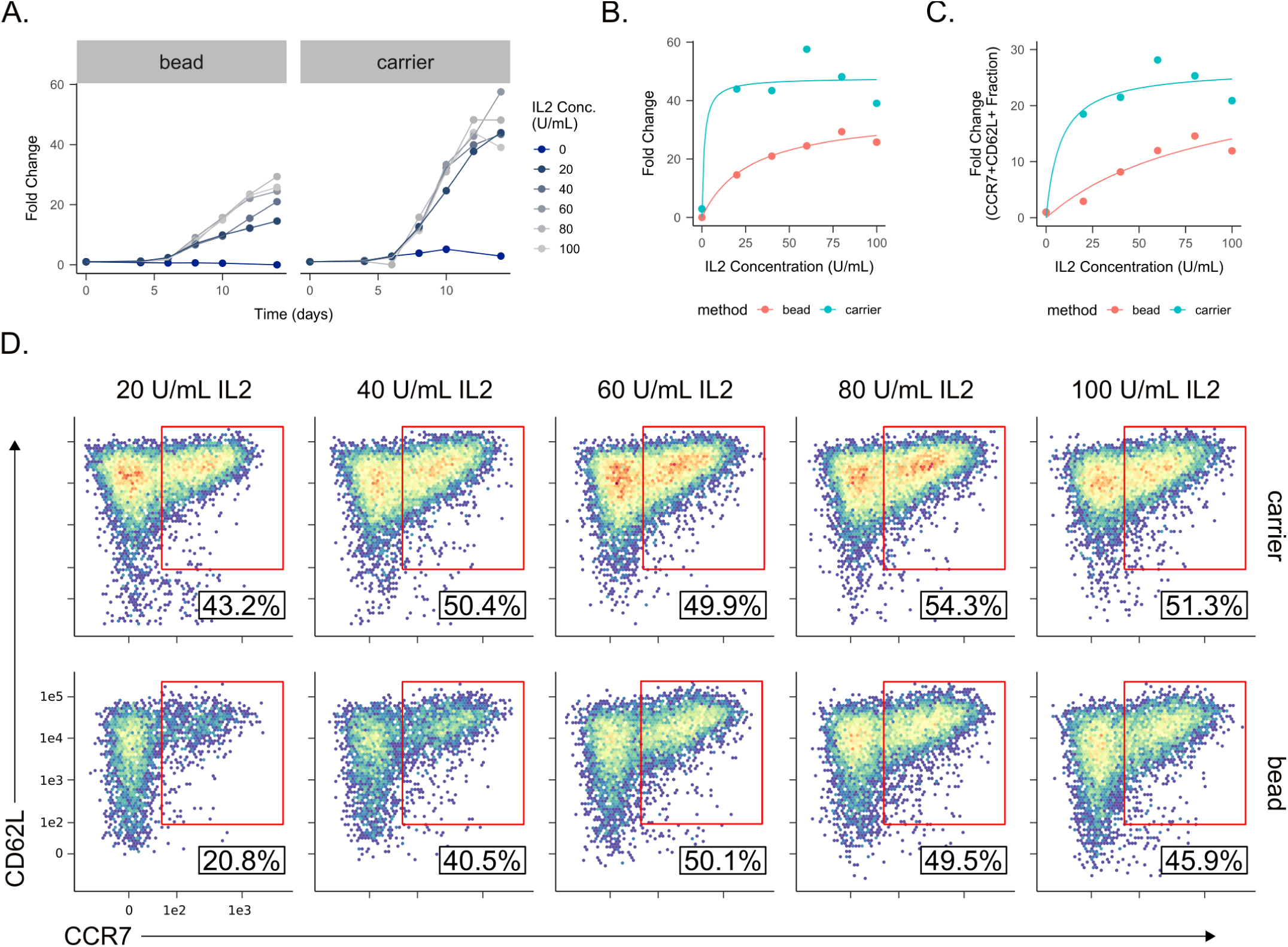
Carriers produce more T cells with less IL2 compared to beads. a) Growth curves of T cells expanded with either beads or carriers grown with varying concentration of IL2. b) Final fold change of T cells grown at varying IL2 concentrations with hyperbolas fitted to plots. c) Final fold change of memory T cells grown at varying IL2 concentration with hyperbolas fitted to plots. d) Flow plots of CC62L+CCR7+ T cells at each IL2 concentration for beads and carriers.

### Microcarriers can be optimized to provide superior memory and CD4 T cell yield

Given that less IL2 was required to expand T cells using the carriers, we sought to optimize the yield of both memory T cells and CD4+ T cells at lower IL2 concentrations. To accomplish this, we employed a design of experiments (DOE) methodology, a technique commonly used to optimize complex manufacturing processes. We varied the IL2 concentration, the number of carriers, and the number of mAbs on the surface of the carriers. Since we desired to understand non-linear influence of these variables, we chose three levels for each (10, 20 and 30 U/mL for IL2; 500, 1500 and 2500 carriers/mL for carrier concentration; 60, 80 and 100 % surface coverage for mAb surface density). Note that in the case of carrier concentration, total cell number was fixed at 2.5 *×* 10^6^ cells/mL, thus this corresponded to seeding densities of 1000, 1666 and 5000 cells/carrier. This led to a randomized 18-run design which included several replicated runs to assess for lack-of-fit (Table S8). T cells were expanded for 14 d using these conditions to modify the expansion process used before.

While there was a wide range of fold changes across all input parameters (Fig. 4a), all runs appeared to generate cells that were greater than 90 % viable when measured using acridine orange/propidium iodide (AO/PI) stain (Fig. 4b). We additionally assessed the percentage of memory and CD4 phenotypes and plotted the number of cells with these markers at day 14. In the case of memory cell yield, IL2 appeared to be highly influential as a main effect, and the other two parameters (carrier concentration and mAb surface density) were less influential (Fig. 4c). Carrier concentration and mAb surface density appeared to have small quadratic effects. For CD4 yield, we noted that all three main effects seem to influence the number of CD4 T cells with little interaction or quadratic effects (Fig. 4d). We further investigated the presence of interaction effects in the memory cell response (Fig. 4e) and noted that there appeared to be interaction between IL2 and carrier concentration (e.g. the slope of one is dependent on the other).

To verify the presence of these qualitative observations in each plot, we produced a model using stepwise linear regression with Akaike information criteria (AIC) as the selection criteria (Table 1 and Figs. S4a and S4b). Neither of these models showed any lack of fit (Tables S9a and S9b), indicating that the generated models accurately described the relationship between the input variables and the response. For memory cell formation, we noted that all main effects were significant. Additionally, we observed significant quadratic effects for carrier concentration and mAb surface density, indicating that these might have an optimum in the middle of the range we tested. Additionally, we found a significant negative interaction effect between carrier concentration and mAb surface density, indicating that there may be antagonism between these two parameters. Using the equation from this model, we calculated the optimum settings for achieving high memory cell yield to be high IL2 (30 U/mL), mid carrier concentration (1500 carriers/mL), and high mAb surface density (approx. 2000 mAbs/µm^2^). For the CD4 response, only the main effects were found significant and all were positively correlated with CD4 cell yield. In this case the optimum settings were simply the high settings for each input (30 U/mL IL2, 2500 carriers/mL, and approx. 2000 mAbs/µm^2^).

**Table 1:** Linear regression output of the DOE experiment for the memory and CD4 T cell yield.

|  | Memory Cells | CD4+ Cells |
| --- | --- | --- |
| Intercept | 2,795,410.000 | -2,375,290.000*** |
| IL2 Conc. | 785,470.300*** | 93,714.670*** |
| Carrier Conc. | 16,050.750*** | 1,528.302*** |
| Ab Density | -24,744.790** | 1,335.737*** |
| (Ab Density)<sup>2</sup> | 10.549*** |  |
| (Carrier Conc.)<sup>2</sup> | -2.584** |  |
| (Ab Dens.)*(Carr. Conc.) | -1.536 |  |
| (IL2 Conc.)*(Carr. Conc.) | -249.904*** |  |
| R<sup>2</sup> | 0.897 | 0.858 |
| Adjusted R<sup>2</sup> | 0.826 | 0.827 |
| *Note:* *p<0.1; **p<0.05; ***p<0.01 |  |  |

**Figure 4:**
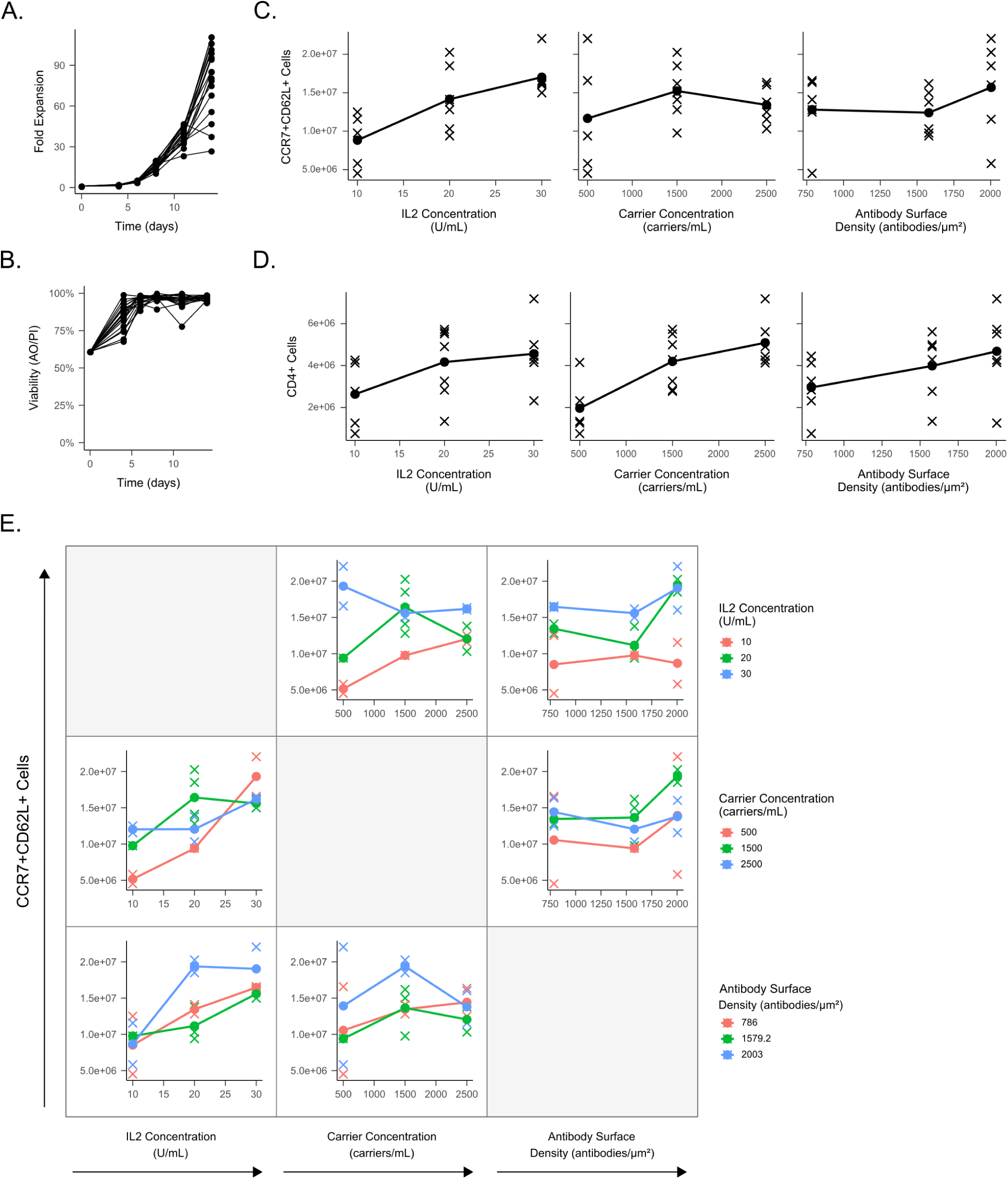
Carriers robustly expand at low IL2 concentrations and have optimal carrier and signal strength inputs. a) Growth curve of carrier expanded cells. b) Viability of carrier expanded cells. c) Main effects plot for memory T cell response. d) Main effects plot for CD4+ T cell response. e) Interaction plots for the DOE with total CCR7+CD62L+ cells as the response. Abbreviations: design of experiments (DOE).

We also performed non-linear symbolic regression analysis to compliment the stepwise linear regression. This was done using the DataModeler software package (Evolved Analytics LLC, Midland, MI) which evolves hundreds to thousands of models using genetic programming and selects the fittest models (those with the highest *R*^2^ and lowest complexity as assessed using a Pareto front) and aggregates them into an ensemble. This has the advantage over linear regression of making fewer *a priori* assumptions about model form. When we fit a model ensemble to memory T cell yield and computed the optimum settings, we observed that the optimal settings were similar to linear regression with the exception of carrier concentration (30 U/mL, 750 carriers/mL and approx. 2000 mAbs/µm^2^) (Fig. 5a). This ensemble consisted of 10 equations which showed robust fit with minimal residual correlations (Figs. S7, S8a, S9a and S10a). When we performed the same analysis for CD4+ T cell yield, we obtained the exact same optimum settings as given by linear regression (30 U/mL IL2, 2500 carriers/mL, and approx. 2000 mAbs/µm^2^). These likewise showed good fit and minimal residual correlation (Figs. S7, S8b, S9b and S10b). Additionally, we plotted the memory and CD4+ T cell yield at the optimal memory settings, and observed a trade-off of the yield between these two subtypes as a function of carrier concentration (Fig. 5c). The optimum setting results from both linear and symbolic regression are summarized in Table 2.

**Table 2:** Summary of predicted process optimums.

| Response | Parameter | Linear Regression | Symbolic Regression |
| --- | --- | --- | --- |
| CCR7+CD62L+ | IL2 Concentration | High | High |
|  | Carrier Concentration | Mid | Low |
|  | mAb Surface Density | High | High |
| CD4+ | IL2 Concentration | High | High |
|  | Carrier Concentration | High | High |
|  | mAb Surface Density | High | High |

**Figure 5:**
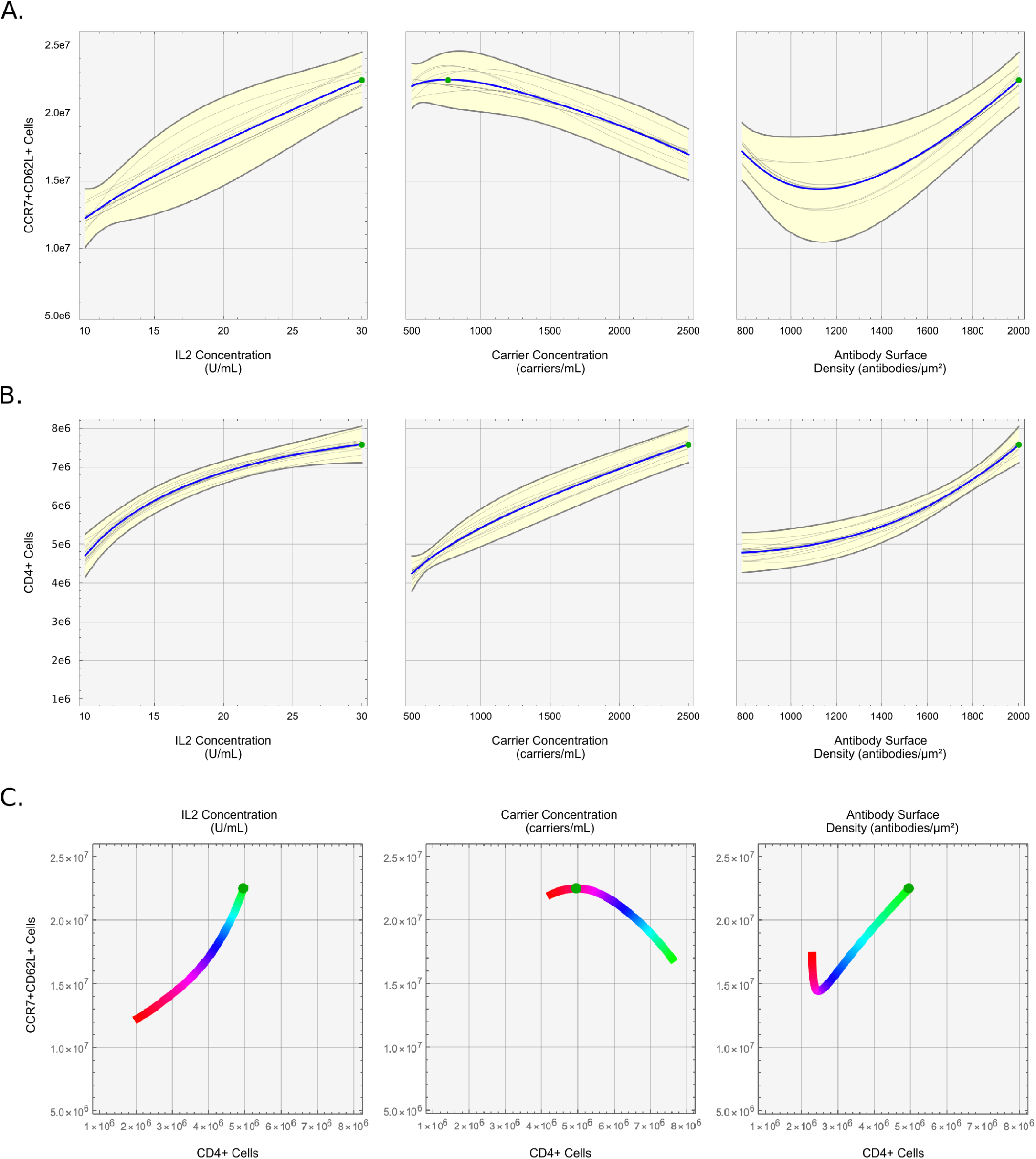
Symbolic regression ensemble plots as given by Evolved Analytics DataModeler. Response profiles of memory (a) and CD4+ (b) vs all three input parameters (IL2 concentration, carrier concentration, and mAb surface density) with optimal settings shown as a green dot. The grey lines are paths from a single given model in the ensemble, and the yellow envelopes represent the variation of the ensemble as a function of each parameter. c) Memory vs CD4+ yield as a function of each input parameter (colored lines, where red is low and green is high) as predicted by the model ensemble for each response. The green point is the optimal setting for memory yield.

Since the total yield of each phenotype was the product of the total cell number and the percentage of that phenotype, we also asked if the total memory and CD4 T cell yields were primarily influenced by bulk T cell expansion or the selective differentiation of a particular phenotype. When performing the same regression on the bulk T cell expansion, memory percentage, or CD4 percentage, we noted that the total live cell response had the same variables in its regression output as the memory yield, indicating that this was likely the main driver of this memory yield composite response (Table S10 and Fig. S5c). However, we also noted that the percentage of memory T cells was negatively affected by increasing carrier concentration (and not by any of the other two variables) (Table S10 and Fig. S5a). In contrast, the CD4 percentage was positively affected by the carrier concentration and the mAb surface density (Table S10 and Fig. S5b). Interestingly, IL2 concentration only affected the bulk expansion. Together, these provided evidence that the differentiation and expansion of memory and CD4 cells were somehow opposed in the carrier system, and the desired balance of CD4 cells and memory cells can be determined by selecting the appropriate carrier concentration.

### Microcarriers can be used to expand functional CAR T cells

After optimizing for memory and CD4 yield, we sought to determine if the carriers were compatible with lentiviral transduction protocols used to generate CAR T cells [26, 27]. We added a 24 h transduction step on day 1 of the 14 d expansion to insert an anti-CD19 CAR [28] and subsequently measured the surface expression of the CAR on day 14 (Figs. 6a and 6b). We noted that there was robust CAR expression in over 25 % of expanded T cells, and there was no observable difference in CAR expression between beads and carriers.

**Figure 6:**
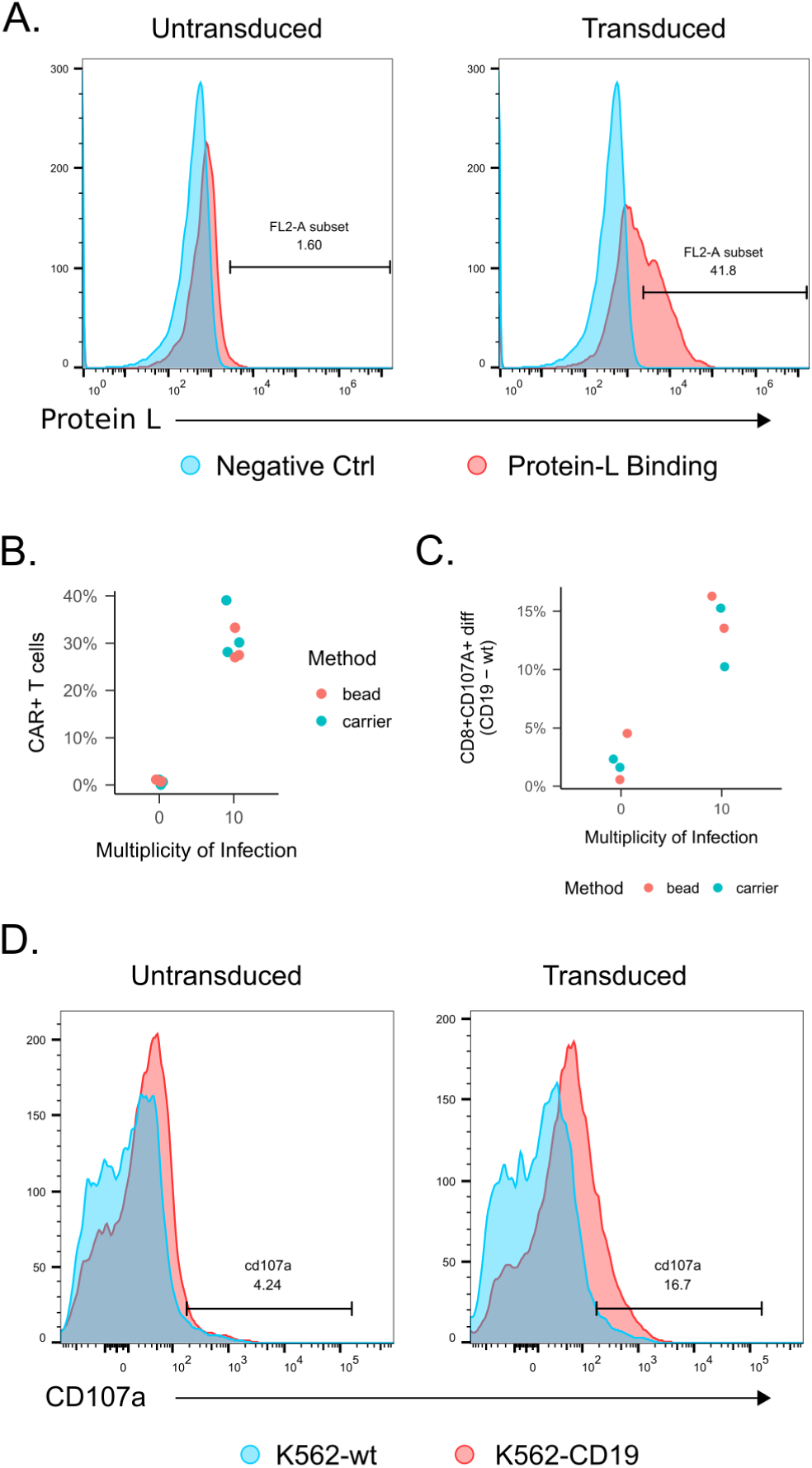
Carriers produce functional CAR T cells. a) CAR expression on day 14 as assessed via flow cytometry for protein L. b) Flow plots from (a) quantified and plotted. c) T cells at day 14 tested for cytotoxicity by measuring CD107a degranulation marker on CD8+ T cells using K562 wild-type or K562-CD19 target cells. Each data point is plotted as a difference in CD107a expression between CD19 and wild-type K562 target cell cultures. d) Flow plots for the degranulation assay shown in (c).

We also verified the functionality of expanded CAR T cells using a degranulation assay [29]. Briefly, T cells were cocultured with target cells (either wild-type K562 or CD19-expressing K562 cells) for 4 h, after which the culture was analyzed via flow cytometry for the appearance of CD107a on CD8+ T cells. CD107a is found on the inner-surface of cytotoxic granules and will emerge on the surface after cytotoxic T cells are activated and degranulate. Indeed, we observed degranulation in T cells expanded with both beads and carriers, although not to an observably different degree (Figs. 6c and 6d). Taken together, these results indicated that the carriers provide similar transduction efficiency compared to beads.

## Discussion

We have developed a T cell expansion system that recapitulates key features of the *in vivo* lymph node microenvironment using microcarriers functionalized with activating mAbs. This strategy provided superior expansion with higher frequency of memory and CD4+ T cells compared to state-of-the-art microbead technology (Fig. 2). Other groups have used biomaterials approaches to mimic the *in vivo* microenvironment [13–15, 17, 30]; however, to our knowledge this is the first system that specifically drives memory and CD4+ T cell formation in a scalable, potentially bioreactor-compatible manufacturing process.

Memory T cells have been shown to be important clinically. Compared to effectors, they have a higher proliferative capacity and are able to engraft for months; thus they are able to provide long-term immunity with smaller doses [19, 31]. Indeed, less differentiated T cells have led to greater survival both in mouse tumor models and human patients [20, 32, 33]. Furthermore, clinical response rates have been positively correlated with T cell expansion, implying that highly-proliferative memory T cells are a significant contributor [18, 34]. Circulating memory T cells have also been found in complete responders who received CAR T cell therapy [35].

Similarly, CD4 T cells have been shown to play an important role in CAR T cell immunotherapy. It has been shown that CAR T doses with only CD4 or a mix of CD4 and CD8 T cells confer greater tumor cytotoxicity than only CD8 T cells [22, 36]. There are several possible reasons for these observations. First, CD4 T cells secrete pro-inflammatory cytokines upon stimulation which may have a synergistic effect on CD8 T cells. Second, CD4 T cells may be less prone to exhaustion and may more readily adopt a memory phenotype compared to CD8 T cells [22]. Third, CD8 T cells may be more susceptible than CD4 T cells to dual stimulation via the CAR and endogenous T Cell Receptor (TCR), which could lead to overstimulation, exhaustion, and apoptosis [23]. Despite evidence for the importance of CD4 T cells, more work is required to determine the precise ratios of CD4 and CD8 T cell subsets to be included in CAR T cell therapy given a disease state.

There are several plausible explanations for the observed phenotypic differences between beads and carriers. First, the carriers are composed of a collagen derivative (gelatin); collagen has been shown to costimulate activated T cells via α1β1 and α2β1 integrins, leading to enhanced proliferation, increased IFNγ production, and upregulated CD25 (IL2Rα) surface expression [8, 10, 11, 37, 38]. Second, there is evidence that providing a larger contact area for T cell activation provides greater stimulation [16, 39]; the carriers have a rougher interface than the 5 µm magnetic beads, and thus could facilitate these larger contact areas. Third, the carriers may allow the T cells to cluster more densely compared to beads, as evidenced by the large clusters on the outside of the carriers (Fig. 1f) as well as the significant fraction of carriers found within their interiors (Figs. S1a and S1b). This may alter the local cytokine environment and trigger different signaling pathways. Particularly, IL15 is secreted by T cells and known to drive memory phenotype [40–42]. The higher cell density in carrier cultures could lead to greater IL15 signaling and thus higher memory cell formation.

An important aspect to our study was the inclusion of a DOE, which is used in many other non-biological disciplines for process development. We specifically used this strategy here to optimize three process variables that plausibly affected T cell growth and phenotype differentiation (IL2 concentration, carrier concentration, and mAb surface density). Additionally, a DOE can facilitate generation of new hypotheses that may explain the potential interactions between parameters. For carrier concentration, we reasoned that this would be directly related to the available surface area, and thus control the degree to which T cell cluster and aggregate. Surprisingly, carrier concentration negatively affected memory cell formation but positively affected CD4 cell formation (Fig. S5 and Table S10). In the case of memory, IL15 is known to drive memory phenotype [40–42], thus the negative relationship between memory fraction and carrier concentration could indicate that other cytokines may start to dominate when the local cell density is increased. In addition to carrier concentration, we varied IL2 concentration as this cytokine has long been known to be required for T cell growth. Our data showed that carriers can lead to robust growth at low IL2 concentrations (20 U/mL) (Fig. 3), thus we decided to investigate this low range in further detail. The DOE revealed that IL2 only affected growth and not phenotype differentiation, as IL2 did not significantly affect memory or CD4 percentage (Fig. S5 and Table S10). This may be because the local IL2 concentration around the carriers was high enough to lessen the effect of exogenous IL2. Furthermore, while the bulk expansion of T cells was influenced by IL2, the negative interaction effect between the IL2 and carrier concentration suggested that IL2 is less influential as the carrier concentration is increased and the cells are clustered more closely, further supporting the hypothesis that the carriers alter the local cytokine concentration. Finally, mAb surface density positively influenced both memory and CD4 T cell formation. This was surprising in the case of memory, since higher stimulation biases differentiation toward effector phenotypes [43]. It could be that in our case, the higher stimulation also increased cytokine output such as IL15, which in turn drove memory cell formation. Ultimately, while the DOE provided an optimal set of conditions that can be used to produce higher numbers of T cells in practice, the surprising findings also generated interesting hypotheses that may lead to better mechanistic understanding and further optimization. These are undergoing further investigation.

In addition to obtaining better phenotypes, other advantages of our carrier approach are that the carriers are large enough to be filtered (approximately 300 µm) using standard 40 µm cell filters or similar. If the remaining cells inside that carriers are also desired, digestion with dispase or collagenase can be used. Furthermore, our system should be compatible with large-scale static culture systems such as the G-Rex bioreactor or perfusion culture systems, which have been previously shown to work well for T cell expansion [12, 44, 45].

It is important to note that all T cell cultures in this study were performed up to 14 days. Others have demonstrated that potent memory T cells may be obtained simply by culturing T cells as little as 5 days using traditional beads [29]. It is unknown if the memory phenotype of our carrier system could be further improved by reducing the culture time, but we can hypothesize that similar results would be observed given the lower number of doublings in a 5 day culture. We should also note that we investigated one memory subtype (CCR7+CD62L+) in this study. Future work will focus on other memory subtypes such as tissue resident memory and stem memory T cells, as well as the impact of using the microcarrier system on the generation of these subtypes.

Another advantage is that the carrier system appears to induce a faster growth rate of T cells given the same IL2 concentration compared to beads (Fig. 3) along with retaining memory phenotype. This has benefits in multiple contexts. Firstly, some patients have small starting T cell populations (such as infants or those who are severely lymphodepleted), and thus require more population doublings to reach a usable dose. Our data suggests the time to reach this dose would be reduced, easing scheduling a reducing cost. Secondly, the allogeneic T cell model would greatly benefit from a system that could create large numbers of T cells with memory phenotype. In contrast to the autologous model which is currently used for *Kymriah* and *Yescarta*, allogeneic T cell therapy would reduce cost by spreading manufacturing expenses across many doses for multiple patients [46]. Since it is economically advantageous to grow as many T cells as possible in one batch in the allogeneic model (reduced start up and harvesting costs, fewer required cell donations), the carriers offer an advantage over current technology.

Finally, while we have demonstrated the carrier system in the context of CAR T cells, this method can theoretically be applied to any T cell immunotherapy which responds to anti-CD3/CD28 mAb and cytokine stimulation. These include tumor infiltrating lymphocytes (TILs), virus-specific T cells (VSTs), T cells engineered to express γδTCR (TEGs), γδ T cells, T cells with transduced-TCR, and CAR-TCR T cells [47–52]. Similar to CD19-CARs used in liquid tumors, these T cell immunotherapies would similarly benefit from the increased proliferative capacity, metabolic fitness, migration, and engraftment potential characteristic of memory phenotypes [53–55]. Indeed, since these T cell immunotherapies are activated and expanded with either soluble mAbs or bead-immobilized mAbs, our system will likely serve as a drop-in substitution to provide these benefits.

## Conclusions

In summary, we have developed an *in vivo*-inspired T cell expansion system using porous, degradable, gelatin microcarriers functionalized with anti-CD3 and anti-CD28 mAbs. Using this system, we have shown that we can achieve higher frequencies of clinically-relevant memory and CD4+ T cell phenotypes compared to traditional bead-based approaches. Additionally, we have shown that they still achieve greater fold change and memory T cell yield beads at low-IL2 concentrations (20 U/mL), and that they can generate functional CAR T cells using lentiviral transduction methods. This system is highly applicable to current T cell manufacturing processes where it may be used to provide higher quality immunotherapies at a reduced reagent cost.

## Methods

### Microcarrier Functionalization

Gelatin microcarriers (CuS or CuG, GE Healthcare) were suspended at 20 mg/mL in 1X phosphate buffered saline (PBS) and autoclaved. All subsequent steps were done aseptically, and all reactions were carried out at 20 mg/mL carriers at room temperature under constant agitation. SNB (Thermo Fisher or Apex Bio) was dissolved at 10 µM in sterile ultrapure water and 7.5 µL_SNB_/mL_PBS_ was added to carrier suspension and allowed to react for 60 min. After washing the carriers three times in sterile PBS, 40 µg/mL STP (Jackson Immunoresearch) was added and allowed to react for 60 min. After the reaction, supernatent was taken for the binding assay, and the carriers were washed twice using sterile PBS. Biotinylated mAbs against human CD3 and CD28 were combined in a 1:1 mass ratio and added to the carriers at 2 µg_mAbs_/mg_carriers_. In the case of the DOE experiment where variable mAb surface density was utilized, the anti-CD3/anti-CD28 mAb mixture was further combined with a biotinylated isotype control to reduce the overall fraction of targeted mAbs (for example the 60 % mAb surface density corresponded to 3 mass parts anti-CD3, 3 mass parts anti-CD8, and 4 mass parts isotype control). mAbs were allowed to bind to the carriers for 60 min. All mAbs were low endotoxin azide free (Biolegend custom, LEAF specification). Carriers were washed in sterile PBS and washed once again in the cell culture media to be used for the T cell expansion. The concentration of the final carrier suspension was found by taking a 50 µL sample, plating in a well, and imaging the entire well. The image was then manually counted to obtain a concentration.

### Microcarrier Quality Control Assays

STP and mAb binding to the carriers was quantified indirectly using a bicinchoninic acid assay (BCA) kit (Thermo Fisher) according to the manufacture’s instructions, with the exception that the standard curve was made with known concentrations of purified STP or IgG instead of bovine serum albumin (BSA). Absorbance at 592 nm was quantified using a Biotek plate reader.

Open biotin binding sites on the carriers after STP coating was quantified indirectly using FITC-biotin (Thermo Fisher). Briefly, 400 pmol/mL FITC-biotin were added to STP-coated carriers and allowed to react for 20 min at room temperature under constant agitation. The supernatant was quantified against a standard curve of FITC-biotin using a Biotek plate reader.

### T Cell Culture

Cryopreserved primary human T cells were either obtained as sorted CD3 subpopulations (Astarte Biotech) or isolated from PBMCs (Zenbio) using a negative selection MACS kit for the CD3 subset (Miltenyi Biotech). T cells were activated using carriers or CD3/CD28 magnetic beads (Miltenyi Biotech). In the case of beads, T cells were activated at the manufacturer recommended cell:bead ratio of 2:1. In the case of carriers, cells were activated using 2500 carriers/cm^2^ unless otherwise noted. Initial cell density was to 2.0 *×* 10^6^ cells/mL to 2.5 *×* 10^6^ cells/mL in a 96 well plate with 300 µL volume. All media was serum-free Cell Therapy Systems OpTmizer (Thermo Fisher) or TexMACS (Miltentyi Biotech) supplemented with 400 U/mL recombinant human IL2 (Peprotech). Cell cultures were expanded for 14 d as counted from the time of initial seeding and activation. Cell counts and viability were assessed using trypan blue or AO/PI and a Countess Automated Cell Counter (Thermo Fisher). Media was added to cultures every 2 d to 3 d depending on media color or a 300 mg/dL minimum glucose threshold. Media glucose was measured using a ChemGlass glucometer.

### Chemotaxis Assay

Migratory function was assayed using a transwell chemotaxis assay as previously described [56]. Briefly, 3 *×* 10^5^ cells were loaded into a transwell plate (5 µm pore size, Corning) with the basolateral chamber loaded with 600 µL media and 0, 250, or 1000 ng/mL CCL21 (Peprotech). The plate was incubated for 4 h after loading, and the basolateral chamber of each transwell was quantified for total cells using countbright beads (Thermo Fisher). The final readout was normalized using the 0 ng/mL concentration as background.

### Degranulation Assay

Cytotoxicity of expanded CAR T cells was assessed using a degranulation assay as previously described [57]. Briefly, 3 *×* 10^5^ T cells were incubated with 1.5 *×* 10^5^ target cells consisting of either K562 wild type cells (ATCC) or CD19-expressing K562 cells transformed with CRISPR (kindly provided by Dr. Yvonne Chen, UCLA) [58]. Cells were seeded in a flat bottom 96 well plate with 1 µg/mL anti-CD49d (eBioscience), 2 µM monensin (eBioscience), and 1 µg/mL anti-CD28 (eBioscience) (all mAbs functional grade) with 250 µL total volume. After 4 h hour incubation at 37 °C, cells were stained for CD3, CD4, and CD107a and analyzed on a BD LSR Fortessa. Readout was calculated as the percent CD107a+ cells of the total CD8 fraction.

### CAR Expression

CAR expression was quantified as previously described [59]. Briefly, cells were washed once and stained with biotinylated Protein L (Thermo Fisher). After a subsequent wash, cells were stained with PE-STP (Biolegend), washed again, and analyzed on a BD Accuri. Readout was percent PE+ cells as compared to secondary controls (PE-STP with no Protein L).

### CAR Plasmid and Lentiviral Transduction

The anti-CD19-CD8-CD137-CD3z chimeric antigen receptor with the EF1α promotor [28] was synthesized (Aldevron) and subcloned into a FUGW lentiviral transfer plasmid (Emory Viral Vector Core). Lentiviral vectors were synthesized by the Emory Viral Vector Core or the Cincinnati Children’s Hospital Medical Center Viral Vector Core.

To transduce primary human T cells, retronectin (Takara) was coated onto non-TC treated 96 well plates and used to immobilize lentiviral vector particles according to the manufacturer’s instructions. Briefly, retronectin solution was adsorbed overnight at 4 °C and blocked the next day using BSA. Prior to transduction, lentiviral supernatant was spinoculated at 2000 *×*g for 2 h at 4 °C. T cells were activated in 96 well plates using beads or carriers for 24 h, and then cells and beads/carriers were transferred onto lentiviral vector coated plates and incubated for another 24 h. Cells and beads/carriers were removed from the retronectin plates using vigorous pipetting and transferred to another 96 well plate wherein expansion continued.

### Statistical Analysis

Statistical significance was evaluated using least-squares linear regression using the *lm* function in R. Stepwise regression models were obtained using the *stepAIC* function from the *MASS* package with forward and reverse stepping. All results with categorical variables are reported relative to baseline reference. Each linear regression was assessed for validity using residual plots (to assess constant variance and independence assumptions), QQ-plots and Shapiro-Wilk normality test (to assess normality assumptions), Box-Cox plots (to assess need for power transformations), and lack-of-fit tests where replicates were present (to assess model fit in the context of pure error). Statistical significance was evaluated at α = 0.05.

For the DOE analysis, the design matrix was created using JMP 13.1 (SAS) with the custom design tool using I-optimal criterion (to minimize prediction variance) and 4 replicates with 2 center points. The experiment was analyzed using linear regression techniques (as described above).

All summary tables were generated using the *stargazer* package in R [60].

### Flow Cytometry Antibodies

All mAbs used for flow cytometry are outlined in Table S11.

### Symbolic Regression

Symbolic regression was done using Evolved Analytics’ DataModeler software. DataModeler uses genetic programming to evolve many symbolic regression models, and then selects the fittest models defined as those with the best trade-off of *R*^2^ (fit) and complexity (this selection accomplished via a pareto front and identifying models at the knee). The collection of fittest models forms a diverse ensemble; the models in the ensemble will agree at observed data points but diverge in extrapolated parameter spaces, providing a trust metric. Feature selection can also be achieved by investigating which variables are present in the fittest models within the ensemble.

In this analysis, DataModeler’s SymbolicRegression function was used to develop nonlinear algebraic models. The fittest models were analyzed to identify dominant variables using the VariablePresence and VariableCombinations functions. CreateModelEnsemble was used to define trustable models using selected variable combinations and these were evaluated using the ModelSummaryTable to identify key statistical attributes with prediction and residual performance assessed visually via the EnsemblePredictionPlot and EnsembleResidualPlot functions, respectively.

Models were developed targeting Memory and CD4 cells with maxima calculated using the ResponsePlotExplorer function. Trade-off performance between these two attributes were explored using the MultiTargetResponseExplorer and ResponseComparisonExplorer with additional insights derived from the ResponseContourPlotExplorer. Graphics and tables were generated by DataModeler.

## Author Contributions

N.J.D., H.W.S., T.K. and K.R. wrote and edited the manuscript. N.J.D., H.W.S, and K.R. designed the experiments. N.J.D. and A.P. optimized the microcarriers and designed the quality control assays. N.J.D. and H.W.S. cultured the T cells and executed the cellular assays. N.J.D. and H.W.S. analyzed the data and generated the figures. T.K. performed symbolic regression. K.R. acquired funding to support the researchers and personnel.

### Acknowledgments

This work was supported by the Cell Manufacturing Technologies (CMaT) National Science Foundation (NSF) Engineering Research Center (Grant EEC-1648035), the NSF Early-Concept Grants for Exploratory Research (EAGER) program (grant 1547638), and the Marcus Foundation, the Georgia Research Alliance, and the Georgia Tech Foundation through their support of the Marcus Center for Therapeutic Cell Characterization and Manufacturing (MC3M) at Georgia Tech. N.D. was supported by the NSF Graduate Research Fellowships Program and the NSF Integrative Graduate Education and Research Traineeship (IGERT, grant 0965945). H.W.S was supported by the NSF EAGER program (grant 1547638). The authors also acknowledge the Viral Vector Core of the Emory Neuroscience National Institute of Neurological Disorders and Stroke (NINDS) Core Facilities (grant P30NS055077) and the Viral Vector Core at Cincinnati Children’s Hospital Medical Center. The authors also thank Dr. Yvonne Chen and Eugenia Zah (UCLA) for providing the K562-CD19+ tumor cells.

## Acronyms

ACT: adoptive cell therapies. 1
AIC: Akaike information criteria. 10
AO/PI: acridine orange/propidium iodide. 10, 17
APC: antigen presenting cell. 2
BCA: bicinchoninic acid assay. 17
BSA: bovine serum albumin. 17, 18
CAR: chimeric antigen receptor. 1, 3, 5, 12–14, 16, 18
CHO: Chinese hamster ovary. 2
CuG: Cultispher-G. 3, 4, 16
CuS: Cultispher-S. 3, 4, 16
DC: dendritic cell. 2
DOE: design of experiments. 1, 7, 8, 14–16, 19
ECM: extracellular matrix. 1, 2
GMP: Good Manufacturing Practices. 2
IgG: immunoglobulin G. 3, 4, 17
mAb: monoclonal antibody. 1–4, 7, 10–19
MACS: magnetic activated cell sorting. 5, 17
MHC: major histocompatibility complex. 2
MTT: 3-(4,5-dimethylthiazol-2-yl)-2,5-diphenyltetrazolium bromide. 3
PBMC: peripheral blood mononuclear cell. 5, 17
PBS: phosphate buffered saline. 16
PDMS: polydimethylsiloxane. 2
SNB: sulfo-NHS-biotin. 3, 4, 16
STP: streptavidin. 3, 4, 16–18
TCR: T Cell Receptor. 14, 16
TEG: T cell engineered to express γdTCR. 16
TIL: tumor infiltrating lymphocyte. 16
VST: virus-specific T cell. 16

## References

[1] Fesnak, A. D., June, C. H. & Levine, B. L. Engineered t cells: the promise and challenges of cancer immunotherapy. Nature Reviews Cancer 16, 566–581 (2016).

[2] Rosenberg, S. A. & Restifo, N. P. Adoptive cell transfer as personalized immunotherapy for human cancer. Science 348, 62–68 (2015).

[3] Roddie, C., O’Reilly, M., Pinto, J. D. A., Vispute, K. & Lowdell, M. Manufacturing chimeric antigen receptor t cells: issues and challenges. Cytotherapy (2019).

[4] Dwarshuis, N. J., Parratt, K., Santiago-Miranda, A. & Roy, K. Cells as advanced therapeutics: State-of-the-art, challenges, and opportunities in large scale biomanufacturing of high-quality cells for adoptive immunotherapies. Advanced Drug Delivery Reviews 114, 222–239 (2017).

[5] Wang, X. & Rivière, I. Clinical manufacturing of CAR t cells: foundation of a promising therapy. Molecular Therapy - Oncolytics 3, 16015 (2016).

[6] Piscopo, N. J. et al. Bioengineering solutions for manufacturing challenges in CAR t cells. Biotechnology Journal 13, 1700095 (2017).

[7] Bashour, K. T. et al. Functional characterization of a t cell stimulation reagent for the production of therapeutic chimeric antigen receptor t cells. Blood 126, 1901–1901 (2015).

[8] Gendron, S., Couture, J. & Aoudjit, F. Integrin *α*2*β*1inhibits fas-mediated apoptosis in t lymphocytes by protein phosphatase 2a-dependent activation of the MAPK/ERK pathway. Journal of Biological Chemistry 278, 48633–48643 (2003).

[9] Ohtani, O. & Ohtani, Y. Structure and function of rat lymph nodes. Archives of Histology and Cytology 71, 69–76 (2008).

[10] Boisvert, M., Gendron, S., Chetoui, N. & Aoudjit, F. Alpha2beta1 integrin signaling augments t cell receptor-dependent production of interferon-gamma in human t cells. Molecular Immunology 44, 3732–3740 (2007).

[11] Ben-Horin, S. & Bank, I. The role of very late antigen-1 in immune-mediated inflammation. Clinical Immunology 113, 119–129 (2004).

[12] Forget, M.-A. et al. Activation and propagation of tumor-infiltrating lymphocytes on clinical-grade designer artificial antigen-presenting cells for adoptive immunotherapy of melanoma. Journal of Immunotherapy 37, 448–460 (2014).

[13] Cheung, A. S., Zhang, D. K. Y., Koshy, S. T. & Mooney, D. J. Scaffolds that mimic antigen-presenting cells enable ex vivo expansion of primary T cells. Nature Biotechnology 36, 160–169 (2018).

[14] del Río, E. P., Miguel, M. M., Veciana, J., Ratera, I. & Guasch, J. Artificial 3d culture systems for t cell expansion. ACS Omega 3, 5273–5280 (2018).

[15] Delalat, B. et al. 3D printed lattices as an activation and expansion platform for T cell therapy. Biomaterials 140, 58–68 (2017).

[16] Meyer, R. A. et al. Immunoengineering: Biodegradable nanoellipsoidal artificial antigen presenting cells for antigen specific t-cell activation (small 13/2015). Small 11, 1612–1612 (2015).

[17] Lambert, L. H. et al. Improving T Cell Expansion with a Soft Touch. Nano letters 17, 821–826 (2017).

[18] Xu, Y. et al. Closely related t-memory stem cells correlate with in vivo expansion of car.cd19-t cells and are preserved by il-7 and il-15. Blood 123, 3750–3759 (2014).

[19] Gattinoni, L., Klebanoff, C. A. & Restifo, N. P. Paths to stemness: building the ultimate antitumour T cell. Nature reviews. Cancer 12, 671–84 (2012).

[20] Fraietta, J. A. et al. Determinants of response and resistance to CD19 chimeric antigen receptor (CAR) t cell therapy of chronic lymphocytic leukemia. Nature Medicine 24, 563–571 (2018).

[21] Gattinoni, L. et al. A human memory t cell subset with stem cell–like properties. Nature Medicine 17, 1290–1297 (2011).

[22] Wang, D. et al. Glioblastoma-targeted CD4+ CAR t cells mediate superior antitumor activity. JCI Insight 3 (2018).

[23] Yang, Y. et al. TCR engagement negatively affects CD8 but not CD4 CAR t cell expansion and leukemic clearance. Science Translational Medicine 9, eaag1209 (2017).

[24] Heathman, T. R. J. et al. Expansion, harvest and cryopreservation of human mesenchymal stem cells in a serum-free microcarrier process. Biotechnology and Bioengineering 112, 1696–1707 (2015).

[25] Sart, S., Errachid, A., Schneider, Y.-J. & Agathos, S. N.Controlled expansion and differentiation of mesenchymal stem cells in a microcarrier based stirred bioreactor. BMC Proceedings 5 (2011).

[26] Tumaini, B. et al. Simplified process for the production of anti-CD19-CAR-engineered T cells. Cytotherapy 15, 1406–15 (2013).

[27] Lamers, C. H. J. et al. T cell receptor-engineered T cells to treat solid tumors: T cell processing toward optimal T cell fitness. Human gene therapy methods 25, 345–57 (2014).

[28] Milone, M. C. et al. Chimeric receptors containing CD137 signal transduction domains mediate enhanced survival of t cells and increased antileukemic efficacy in vivo. Molecular Therapy 17, 1453–1464 (2009).

[29] Ghassemi, S. et al. Reducing Ex Vivo Culture improves the antileukemic activity of chimeric antigen receptor (CAR) t cells. Cancer Immunology Research 6, 1100–1109 (2018).

[30] Matic, J., Deeg, J., Scheffold, A., Goldstein, I. & Spatz, J. P. Fine tuning and efficient T cell activation with stimulatory aCD3 nanoarrays. Nano letters 13, 5090–7 (2013).

[31] Joshi, N. S. & Kaech, S. M. Effector CD8 t cell development: A balancing act between memory cell potential and terminal differentiation. The Journal of Immunology 180, 1309–1315 (2008).

[32] Adachi, K. et al. IL-7 and CCL19 expression in CAR-t cells improves immune cell infiltration and CAR-t cell survival in the tumor. Nature Biotechnology 36, 346–351 (2018).

[33] Rosenberg, S. A. et al. Durable complete responses in heavily pretreated patients with metastatic melanoma using t-cell transfer immunotherapy. Clinical Cancer Research 17, 4550–4557 (2011).

[34] Besser, M. J. et al. Clinical responses in a phase II study using adoptive transfer of short-term cultured tumor infiltration lymphocytes in metastatic melanoma patients. Clinical Cancer Research 16, 2646–2655 (2010).

[35] Kalos, M. et al. T cells with chimeric antigen receptors have potent antitumor effects and can establish memory in patients with advanced leukemia. Science translational medicine 3, 95ra73 (2011).

[36] Sommermeyer, D. et al. Chimeric antigen receptor-modified t cells derived from defined CD8+ and CD4+ subsets confer superior antitumor reactivity in vivo. Leukemia 30, 492–500 (2015).

[37] Rao, W. H., Hales, J. M. & Camp, R. D. R. Potent costimulation of effector t lymphocytes by human collagen type i. The Journal of Immunology 165, 4935–4940 (2000).

[38] Bank, I., Book, M. & Ware, R. Functional role of VLA-1 (CD49a) in adhesion, cation-dependent spreading, and activation of cultured human t lymphocytes. Cellular Immunology 156, 424–437 (1994).

[39] Hickey, J. W., Vicente, F. P., Howard, G. P., Mao, H.-Q. & Schneck, J. P. Biologically inspired design of nanoparticle artificial antigen-presenting cells for immunomodulation. Nano Letters 17, 7045–7054 (2017).

[40] Gomez-Eerland, R. et al. Manufacture of gene-modified human t-cells with a memory stem/central memory phenotype. Human Gene Therapy Methods 25, 277–287 (2014).

[41] Buck, M. D. et al. Mitochondrial Dynamics Controls T Cell Fate through Metabolic Programming. Cell 166, 114 (2016).

[42] van der Windt, G. J. et al. Mitochondrial respiratory capacity is a critical regulator of CD8+ t cell memory development. Immunity 36, 68–78 (2012).

[43] D’Souza, W. N. & Hedrick, S. M. Cutting edge: Latecomer CD8 t cells are imprinted with a unique differentiation program. The Journal of Immunology 177, 777–781 (2006).

[44] Gerdemann, U., Vera, J. F., Rooney, C. M. & Leen, A. M. Generation of multivirus-specific T cells to prevent/treat viral infections after allogeneic hematopoietic stem cell transplant. Journal of visualized experiments: JoVE (2011).

[45] Jin, J. et al. Simplified method of the growth of human tumor infiltrating lymphocytes in gas-permeable flasks to numbers needed for patient treatment. Journal of immunotherapy (Hagerstown, Md.: 1997) 35, 283–92 (2012).

[46] Harrison, R. P., Zylberberg, E., Ellison, S. & Levine, B. L. Chimeric antigen receptor–t cell therapy manufacturing: modelling the effect of offshore production on aggregate cost of goods. Cytotherapy (2019).

[47] Cho, H.-W. et al. Triple costimulation via CD80, 4-1bb, and CD83 ligand elicits the long-term growth of vγ9vδ2 t cells in low levels of IL-2. Journal of Leukocyte Biology 99, 521–529 (2015).

[48] Straetemans, T. et al. GMP-grade manufacturing of t cells engineered to express a defined γδTCR. Frontiers in Immunology 9 (2018).

[49] Robbins, P. F. et al. Tumor regression in patients with metastatic synovial cell sarcoma and melanoma using genetically engineered lymphocytes reactive with NY-ESO-1. Journal of Clinical Oncology 29, 917–924 (2011).

[50] Brimnes, M. K. et al. Generation of autologous tumor-specific t cells for adoptive transfer based on vaccination, in vitro restimulation and CD3/CD28 dynabead-induced t cell expansion. Cancer Immunology, Immunotherapy 61, 1221–1231 (2012).

[51] Baldan, V., Griffiths, R., Hawkins, R. E. & Gilham, D. E. Efficient and reproducible generation of tumour-infiltrating lymphocytes for renal cell carcinoma. British Journal of Cancer 112, 1510–1518 (2015).

[52] Walseng, E. et al. A TCR-based chimeric antigen receptor. Scientific Reports 7(2017).

[53] Blanc, C. et al. Targeting resident memory t cells for cancer immunotherapy. Frontiers in Immunology 9(2018).

[54] Lalor, S. J. & McLoughlin, R. M. Memory γδ t cells–newly appreciated protagonists in infection and immunity. Trends in Immunology 37, 690–702 (2016).

[55] Rosato, P. C. et al. Virus-specific memory t cells populate tumors and can be repurposed for tumor immunotherapy. Nature Communications 10 (2019).

[56] Hromas, R. et al. Cloning and characterization of exodus, a novel beta-chemokine. Blood 89, 3315–3322 (1997).

[57] Schmoldt, A., Benthe, H. F. & Haberland, G. Digitoxin metabolism by rat liver microsomes. Biochemical pharmacology 24, 1639–1641 (1975).

[58] Zah, E., Lin, M.-Y., Silva-Benedict, A., Jensen, M. C. & Chen, Y. Y. T cells expressing cd19/cd20 bispecific chimeric antigen receptors prevent antigen escape by malignant b cells. Cancer immunology research 4, 498–508 (2016).

[59] Zheng, Z., Chinnasamy, N. & Morgan, R. A. Protein l: a novel reagent for the detection of chimeric antigen receptor (CAR) expression by flow cytometry. Journal of Translational Medicine 10, 29 (2012).

[60] Hlavac, M. *stargazer: Well-Formatted Regression and Summary Statistics Tables*. Central European Labour Studies Institute (CELSI), Bratislava, Slovakia (2018). R package version 5.2.2.

