## Supplemental Tables and figures for "Functionalized microcarriers improve T cell manufacturing by facilitating migratory memory T cell production and increasing CD4/CD8 ratio"

### Supplemental Materials

Table S1: Linear regression output of the cell fraction remaining inside carriers.

| | Carrier Fraction (%) |
|---|---|
| Intercept | 0.136* |
| Initial Cell Number | 0.00000 |
| $R^2$ | 0.186 |
| Adjusted $R^2$ | −0.086 |
| *Note:* | *p<0.1; **p<0.05; ***p<0.01 |

Table S2: Linear regression output of the combined T cell expansion T cell datasets for memory response.

| | Total Memory Percent |
|---|---|
| Intercept | 0.319*** (0.059) |
| Method (Carrier) | 0.128*** (0.025) |
| Donor (2) | −0.106*** (0.030) |
| Donor (3) | −0.516*** (0.102) |
| Donor (4) | −0.640*** (0.098) |
| Donor (5) | −0.214** (0.099) |
| Total Glucose Added (mg) | 0.002 (0.002) |
| Volumetric Fold Change | 0.001 (0.001) |
| Total IL-2 Added (U) | 0.00001 (0.00001) |
| $R^2$ | 0.790 |
| Adjusted $R^2$ | 0.758 |
| *Note:* | *p<0.1; **p<0.05; ***p<0.01 |

Table S3: Linear regression output of the combined T cell expansion T cell datasets for CD4+ response.

| | CD4+ Percent |
|---|---|
| Intercept | −1.971*** (0.222) |
| Method (Carrier) | 0.923*** (0.095) |
| Donor (2) | −0.565*** (0.113) |
| Donor (3) | 0.134 (0.386) |
| Donor (4) | −0.827** (0.369) |
| Donor (5) | 0.013 (0.375) |
| Total Glucose Added (mg) | −0.015** (0.007) |
| Volumetric Fold Change | 0.005 (0.004) |
| Total IL-2 Added (U) | 0.0001* (0.00004) |
| $R^2$ | 0.814 |
| Adjusted $R^2$ | 0.786 |
| *Note:* | *p<0.1; **p<0.05; ***p<0.01 |

Table S4: Linear regression output of the combined T cell expansion T cell datasets for CD4+CCR7+CD62L+ response.

| | log(CD4 Memory Percent) |
|---|---|
| Intercept | −2.343*** (0.238) |
| Method (Carrier) | 1.149*** (0.102) |
| Donor (2) | −0.621*** (0.121) |
| Donor (3) | −0.991** (0.414) |
| Donor (4) | −2.299*** (0.395) |
| Donor (5) | −0.654 (0.402) |
| Total Glucose Added (mg) | −0.026*** (0.007) |
| Volumetric Fold Change | 0.009** (0.004) |
| Total IL2 Added (U) | 0.0001*** (0.00005) |
| $R^2$ | 0.891 |
| Adjusted $R^2$ | 0.875 |
| *Note:* | *p<0.1; **p<0.05; ***p<0.01 |

Table S5: Linear regression output of the combined T cell expansion T cell datasets for CD8+CCR7+CD62L+ response.

| | CD8 Memory Percent |
|---|---|
| Intercept | 0.203*** (0.054) |
| Method (Carrier) | 0.057** (0.023) |
| Donor (2) | −0.065** (0.027) |
| Donor (3) | −0.462*** (0.094) |
| Donor (4) | −0.568*** (0.089) |
| Donor (5) | −0.215** (0.091) |
| Total Glucose Added (md) | 0.004** (0.002) |
| Volumetric Fold Change | 0.001 (0.001) |
| Total IL2 Added (U) | 0.00001 (0.00001) |
| $R^2$ | 0.746 |
| Adjusted $R^2$ | 0.708 |
| *Note:* | *p<0.1; **p<0.05; ***p<0.01 |

Table S6: Linear regression output for CCL21 transwell experiment with percent migration as response.

| | Normalized Migration (%) |
|---|---|
| Intercept | 0.040*** |
| CCL21 Conc. | 0.0001*** |
| Method (Carrier) | 0.051*** |
| MOI | −0.001 |
| $R^2$ | 0.706 |
| Adjusted $R^2$ | 0.662 |
| *Note:* | *p<0.1; **p<0.05; ***p<0.01 |

Table S7: T cell/PBMC donor data.

| ID | Vendor | Age | Demographic | Gender | BMI | CD3+ | CD4+ | CD8+ | CD19+ | CD14+ |
|---|---|---|---|---|---|---|---|---|---|---|
| 1 | Astarte | 27 | Caucasian | Female | 33.4 | 99.0 | 62.5 | 33.6 | 0.1 | 0.3 |
| 2 | Astarte | 23 | Caucasian | Male | 51.4 | 99.1 | 65.9 | 35.7 | 0.3 | 2.0 |
| 3 | Zenbio | 27 | African American | Male | 21.7 | 47.0 | 32.7 | 5.5 | 10.8 | 12.3 |
| 4 | Zenbio | 25 | African American | Male | 20.9 | 57.0 | 35.2 | 18.2 | 9.0 | 15.1 |
| 5 | Zenbio | 28 | African American | Male | 22.9 | 62.9 | 46.5 | 11.9 | 3.1 | 17.2 |

Table S8: Experimental run matrix for the DOE experiment.

| Run | IL2 Conc. (U/mL) | Carrier Conc. (carriers/mL) | Ab Density (abs/$\mu m^2$) |
|---|---|---|---|
| 1 | 30 | 500 | 2003 |
| 2 | 20 | 2500 | 1579.2 |
| 3 | 20 | 1500 | 2003 |
| 4 | 20 | 1500 | 786 |
| 5 | 20 | 1500 | 2003 |
| 6 | 30 | 2500 | 2003 |
| 7 | 10 | 2500 | 786 |
| 8 | 20 | 2500 | 1579.2 |
| 9 | 20 | 500 | 1579.2 |
| 10 | 10 | 2500 | 2003 |
| 11 | 30 | 1500 | 1579.2 |
| 12 | 10 | 1500 | 1579.2 |
| 13 | 30 | 1500 | 1579.2 |
| 14 | 10 | 500 | 786 |
| 15 | 30 | 500 | 786 |
| 16 | 10 | 500 | 2003 |
| 17 | 20 | 1500 | 786 |
| 18 | 30 | 2500 | 786 |

Table S9: ANOVA with lack of fit of the DOE experiment for both memory and CD4 responses.

(a) memory

| | Df | F value | Pr(>F) |
|---|---|---|---|
| Ab Dens. | 1 | 8.013 | 0.047 |
| IL2 Conc. | 1 | 77.541 | 0.001 |
| Carr. Conc. | 1 | 2.902 | 0.164 |
| $(\text{Ab Dens.})^2$ | 1 | 16.439 | 0.015 |
| $(\text{Carr. Conc.})^2$ | 1 | 12.207 | 0.025 |
| (Ab Dens.)*(Carr. Conc.) | 1 | 3.114 | 0.152 |
| (IL2 Conc.)*(Carr. Conc.) | 1 | 21.744 | 0.010 |
| Residuals | 10 | | |
| Lack of fit | 6 | 2.039 | 0.256 |
| Pure Error | 4 | | |

(b) CD4+

| | Df | F value | Pr(>F) |
|---|---|---|---|
| IL2 Conc. | 1 | 63.642 | 0.001 |
| Carr. Conc. | 1 | 169.968 | 0.0002 |
| Ab Dens. | 1 | 53.240 | 0.002 |
| Residuals | 14 | | |
| Lack of fit | 10 | 4.356 | 0.084 |
| Pure Error | 4 | | |

Table S10: Linear regression output of the DOE experiment for each decomposed response (fold change, memory cell percent, and CD4+ cell percent).

| | Memory % | CD4+ % | Total Live Cells |
|---|---|---|---|
| | (1) | (2) | (3) |
| Intercept | 0.452*** | 0.013 | 2,795,410.000 |
| IL2 Conc. | 0.001 | 0.00003 | 785,470.300*** |
| Carrier Conc. | −0.00003** | 0.00004*** | 16,050.750*** |
| Ab Density | 0.00003 | 0.00004*** | −24,744.790** |
| $(\text{Ab Density})^2$ | | | 10.549*** |
| $(\text{Carrier Conc})^2$ | | | −2.584** |
| (IL2 Conc.)*(Carrier Conc.) | | | −249.904*** |
| (Ab Density)*(Carrier Conc.) | | | −1.536 |
| Observations | 18 | 18 | 18 |
| $R^2$ | 0.418 | 0.910 | 0.897 |
| Adjusted $R^2$ | 0.294 | 0.891 | 0.826 |
| Residual Std. Error | 0.041 (df = 14) | 0.013 (df = 14) | 1,931,344.000 (df = 10) |
| F Statistic | 3.355** (df = 3; 14) | 47.457*** (df = 3; 14) | 12.492*** (df = 7; 10) |

*Note:* $^{*}p<0.1$; $^{**}p<0.05$; $^{***}p<0.01$

Table S11: Antibodies used.

| Antigen | Fluorophore | Vendor | Catalog Number | Figure Used |
|---|---|---|---|---|
| CD3 | APC-Fire | Biolegend | 34839 | Figs. 3 and 4 |
| CD3 | APC-H7 | Becton Dickenson | 561438 | Figs. 2 and 6 |
| CD4 | PerCP-Cy5.5 | Biolegend | 344607 | Figs. 3 and 4 |
| CD4 | PerCP-Cy5.5 | Becton Dickenson | 561438 | Figs. 2 and 6 |
| CCR7 | AF647 | Becton Dickenson | 560816 | Figs. 3 and 4 |
| CCR7 | AF647 | Becton Dickenson | 561438 | Fig. 2 |
| CD62L | PE | Becton Dickenson | 341012 | Figs. 3 and 4 |
| CD107a | AF647 | Becton Dickenson | 562622 | Fig. 6 |

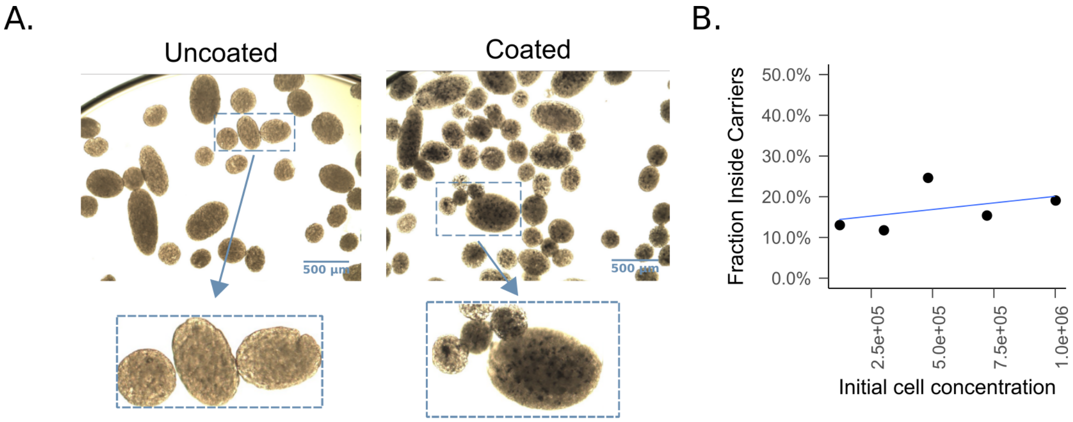


Figure S1: a) T cells inside the carriers as quantified using an automated cell counter after digestion. b) Images of T cells inside carriers as seen using an 3-(4,5-dimethylthiazol-2-yl)-2,5-diphenyltetrazolium bromide (MTT) stain after washing away unbound cells from carriers.

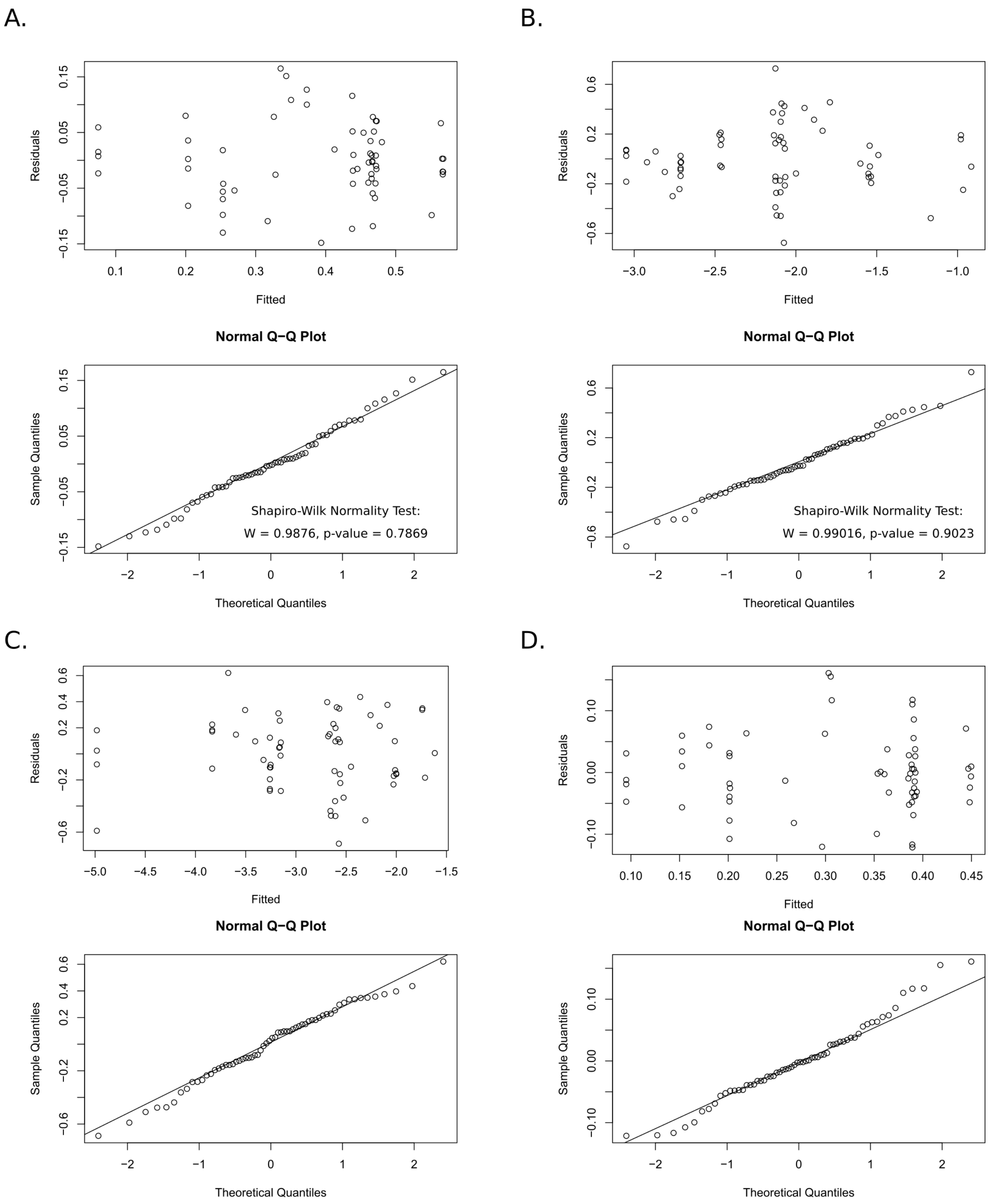


Figure S2: Regression diagnostics including residual plots, Q-Q Normality Plots of the residuals, and the Shapiro-Wilk Test for normality of the residuals for a) memory, b) CD4+ T cells, c). CD4+ memory T cells, and d) CD8+ memory T cells.

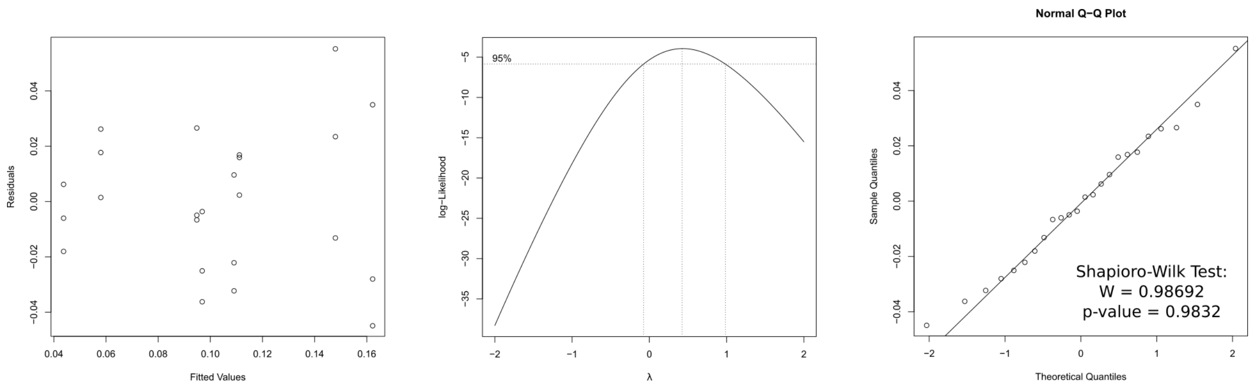


Figure S3: Regression diagnostics for the experiment (Table S6) including residual plots, Q-Q Normality Plots of the residuals, and the Shapiro-Wilk Test for normality of the residuals.

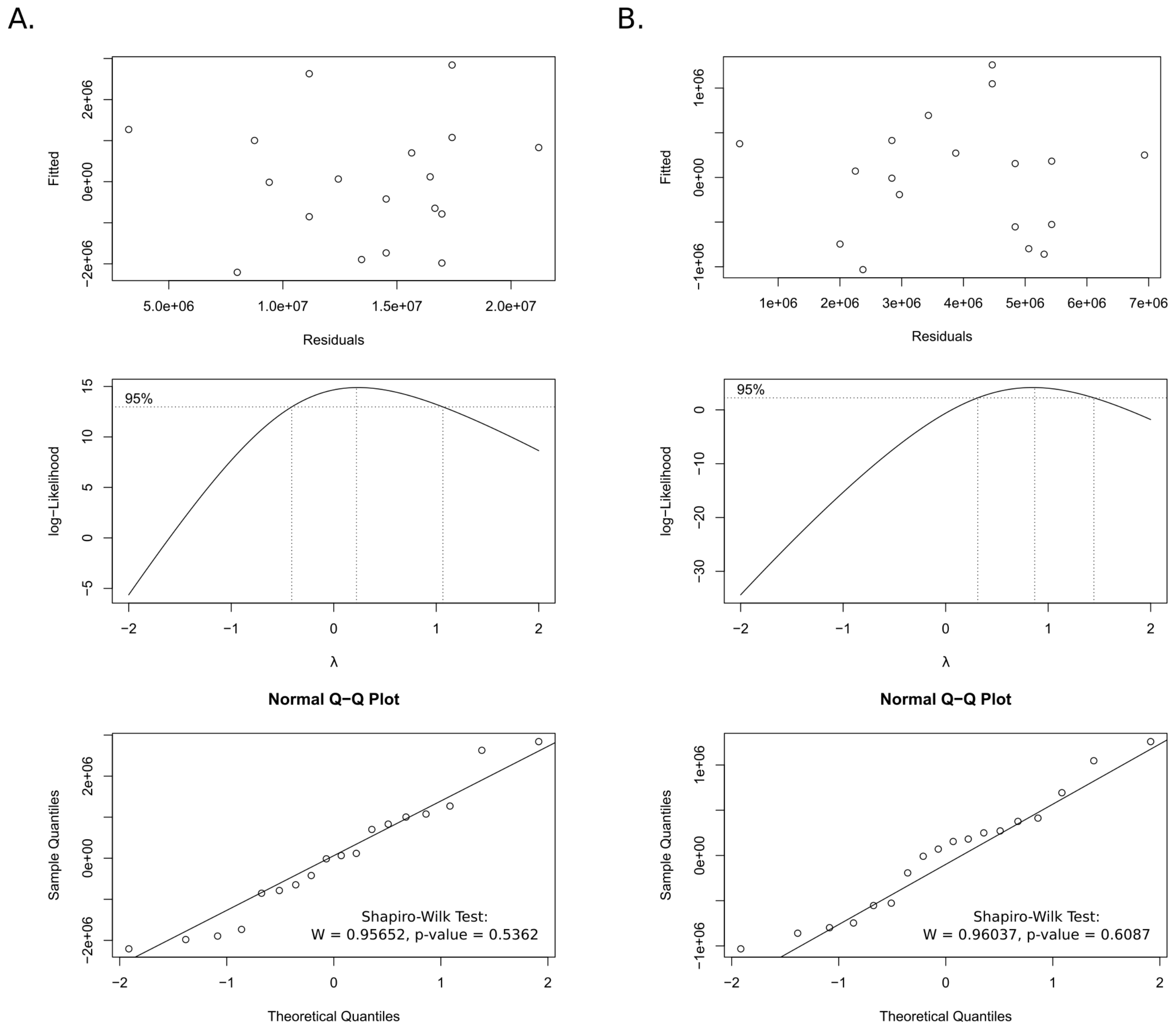


Figure S4: Regression diagnostics including residual plots, Q-Q Normality Plots of the residuals, and the Shapiro-Wilk Test for normality of the residuals for a) memory and b) CD4+ T cells.

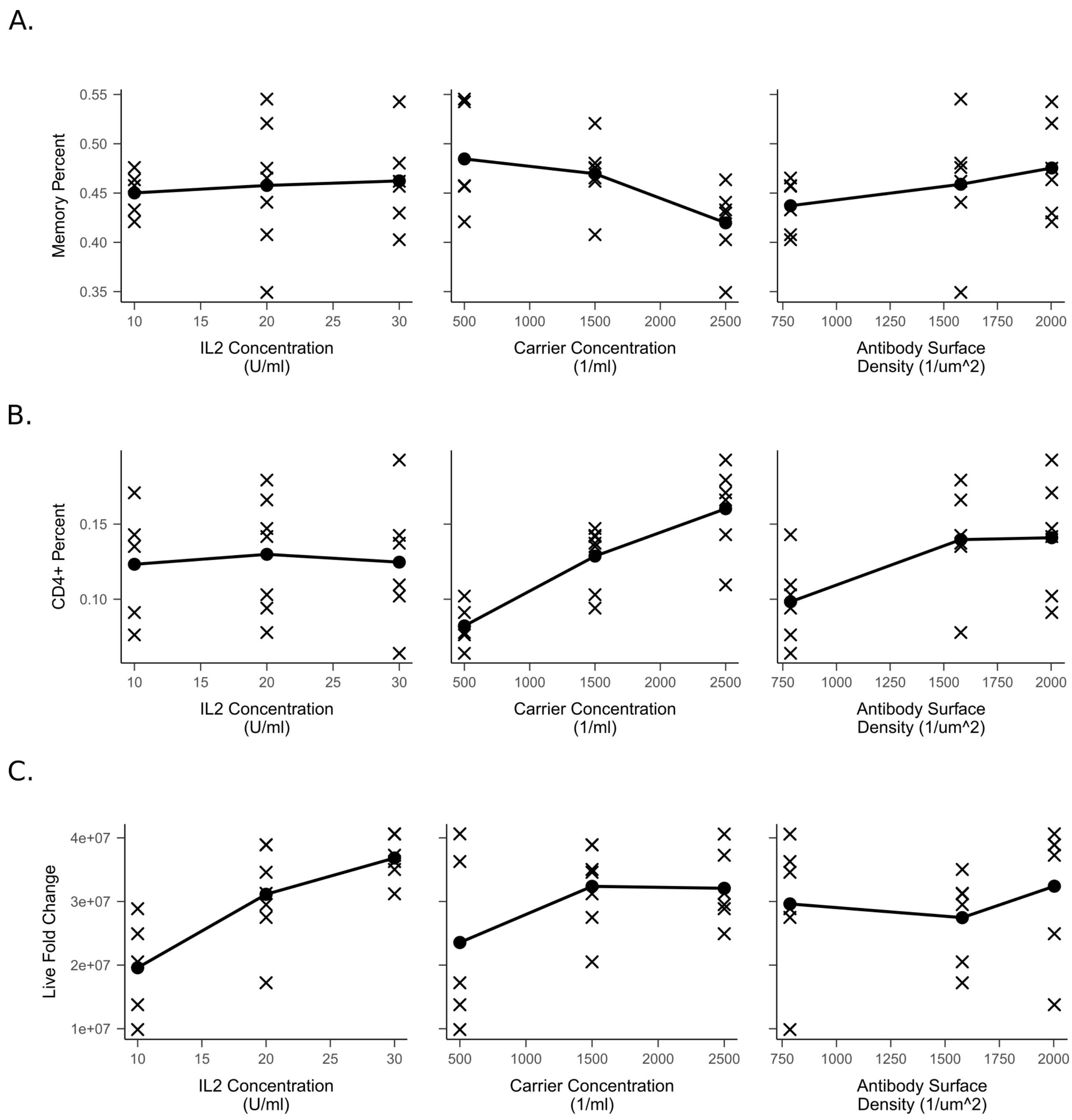


Figure S5: Main effects plots for a) memory percentage, b) CD4+ percentage. and c) total fold change of all cells.

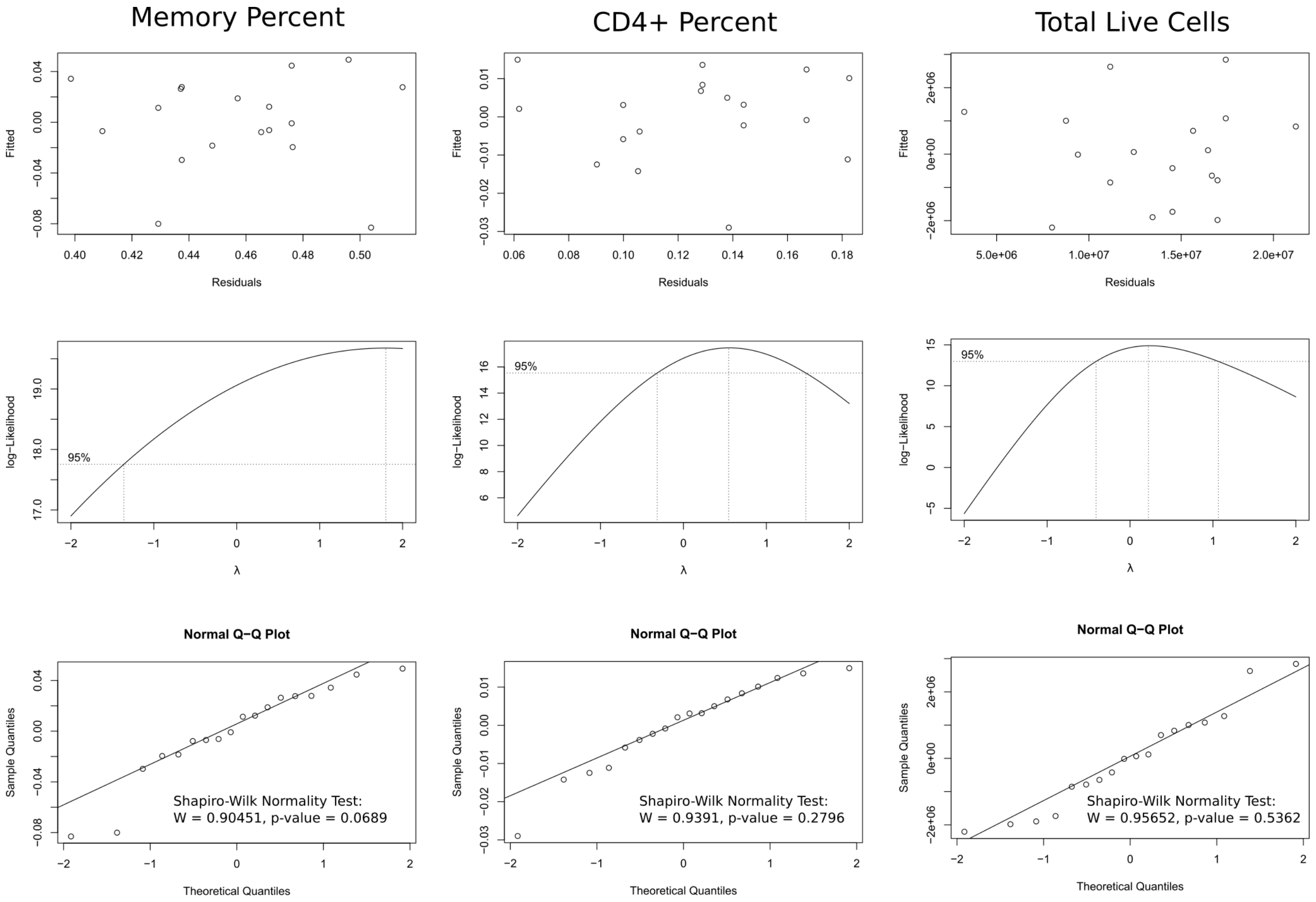


Figure S6: Diagnostics for the decomposed responses in Figs. S5a to S5c.

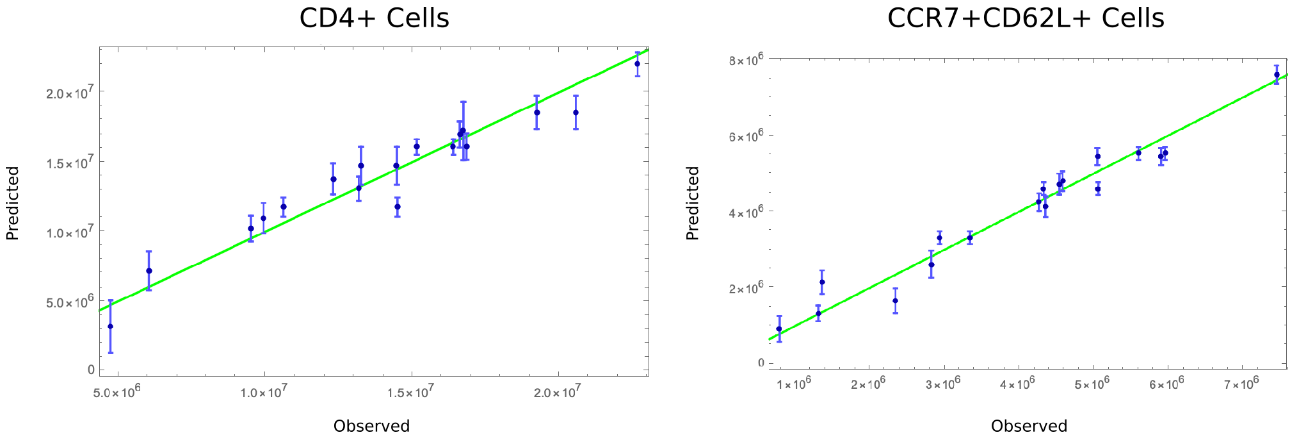


Figure S7: Predicted vs observed plots for symbolic regression analysis.

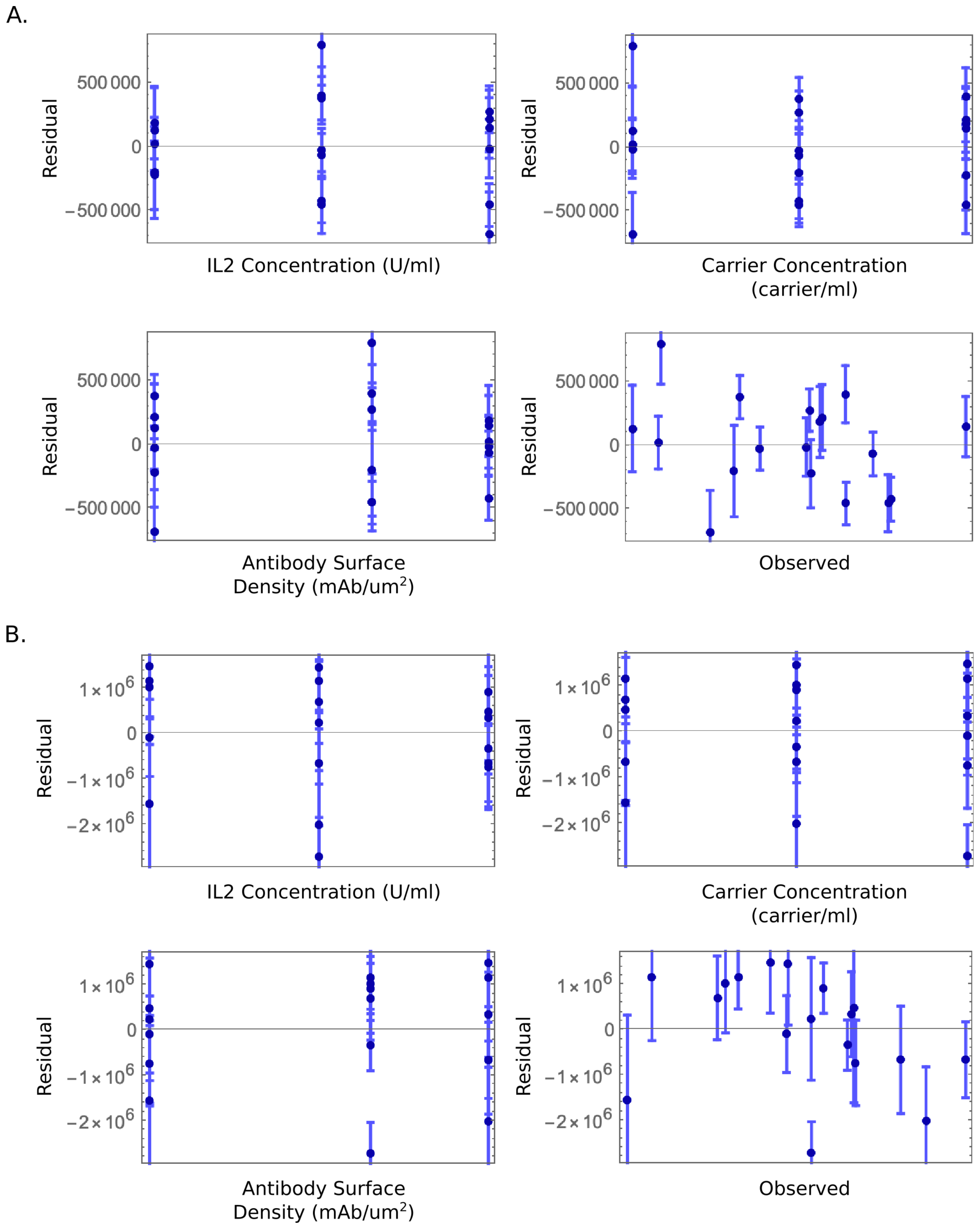


Figure S8: Symbolic regression residuals vs IL2 concentration, carrier concentration, antibody surface density, and the observed response (upper left, upper right, lower left, and lower right panel respectively) for a) memory and b) CD4+ T cell yield responses.

A.

| | | | |
|---|---|---|---|
| Ensemble Quality | $R^2$ | | 0.933507 |
| | # Variables | | 3 |
| | # Models | | 10 |
| | Avg Model Complexity | | 79.4 |
| Model Quality | Model | Complexity | $1-R^2$ |
| | 1 | 61 | 0.0873975 |
| | 2 | 65 | 0.0994461 |
| | 3 | 69 | 0.0873944 |
| | 4 | 82 | 0.0820153 |
| | 5 | 82 | 0.0820765 |
| | 6 | 83 | 0.0819227 |
| | 7 | 83 | 0.0961709 |
| | 8 | 84 | 0.0800521 |
| | 9 | 89 | 0.087623 |
| | 10 | 96 | 0.0794506 |
| Variables | il2•conc | | activator•conc |
| | abs•surface•density | | |
| R–Squared | 0.933507 | | |
| Adjusted R–Squared | 0.929351 | | |
| Noise Power | 0.0064263 | | |

B.

| | | | |
|---|---|---|---|
| Ensemble Quality | $R^2$ | | 0.958883 |
| | # Variables | | 3 |
| | # Models | | 13 |
| | Avg Model Complexity | | 52.6154 |
| Model Quality | Model | Complexity | $1-R^2$ |
| | 1 | 38 | 0.060805 |
| | 2 | 46 | 0.0461456 |
| | 3 | 46 | 0.057544 |
| | 4 | 46 | 0.0597898 |
| | 5 | 49 | 0.0562038 |
| | 6 | 50 | 0.0449207 |
| | 7 | 50 | 0.045076 |
| | 8 | 53 | 0.0419094 |
| | 9 | 54 | 0.045406 |
| | 10 | 55 | 0.0494997 |
| | 11 | 61 | 0.0447944 |
| | 12 | 62 | 0.0482701 |
| | 13 | 74 | 0.0385888 |
| Variables | il2•conc | | activator•conc |
| | abs•surface•density | | |
| R–Squared | 0.958883 | | |
| Adjusted R–Squared | 0.956313 | | |
| Noise Power | 0.00663597 | | |

Figure S9: Symbolic regression model summaries for a) memory and b) CD4+ T cell yield responses. NOTE: il2·conc = IL2 Concentration (U/ml), activator·conc = Carrier Concentration (carriers/ml), abs·surface·density = Antibody Surface Density (mAbs/$\mu m^2$)

A.

$$-\left(4.10\times10^{8}\right)+\frac{2.57\times10^{10}}{\text{abs·surface·density}}+19953.68\,\text{abs·surface·density}-\left(3.92\times10^{-4}\right)\text{abs·surface·density}\,\text{activator·conc}^{2}+\frac{\left(3.79\times10^{8}\right)(-29.01+\text{activator·conc}+\text{il2·conc})}{\text{activator·conc}}$$

$$-40049653.00+\frac{3.00\times10^{10}}{\text{abs·surface·density}}+28970.84\,\text{abs·surface·density}-2.82\,\text{abs·surface·density}\,\text{activator·conc}-\frac{\left(3.58\times10^{9}\right)\left(-8.96+\sqrt{\text{abs·surface·density}}\right)}{\text{activator·conc}\,\text{il2·conc}}$$

$$-\left(4.10\times10^{8}\right)+20093.66\,\text{abs·surface·density}+\frac{2.65\times10^{10}}{13.60+\text{abs·surface·density}}-\left(3.90\times10^{-4}\right)\text{abs·surface·density}\,\text{activator·conc}^{2}+\frac{\left(3.79\times10^{8}\right)(-29.01+\text{activator·conc}+\text{il2·conc})}{\text{activator·conc}}$$

$$-\left(4.00\times10^{8}\right)+\frac{2.64\times10^{10}}{\text{abs·surface·density}}+20624.74\,\text{abs·surface·density}-\left(4.76\times10^{-4}\right)\text{abs·surface·density}\,\text{activator·conc}^{2}+\frac{\left(3.69\times10^{8}\right)(-29.01+\text{activator·conc}+\text{il2·conc})}{2+\text{activator·conc}+\frac{1}{\text{il2·conc}}}$$

$$-\left(4.01\times10^{8}\right)+\frac{2.63\times10^{10}}{\text{abs·surface·density}}+20609.07\,\text{abs·surface·density}-\left(4.76\times10^{-4}\right)\text{abs·surface·density}\,\text{activator·conc}^{2}+\frac{\left(3.70\times10^{8}\right)(-29.01+\text{activator·conc}+\text{il2·conc})}{2+\frac{1}{\text{abs·surface·density}}+\text{activator·conc}}$$

$$-\left(3.99\times10^{8}\right)+\frac{2.65\times10^{10}}{\text{abs·surface·density}}+20757.62\,\text{abs·surface·density}-\left(4.97\times10^{-4}\right)\text{abs·surface·density}\,\text{activator·conc}^{2}+\frac{\left(3.67\times10^{8}\right)(-29.01+\text{activator·conc}+\text{il2·conc})}{\frac{5}{2}+\text{activator·conc}}$$

$$-29156378.00+\frac{2.96\times10^{10}}{\text{abs·surface·density}}+23528.24\,\text{abs·surface·density}-\left(7.72\times10^{-4}\right)\text{activator·conc}\left(-2+\sqrt{\text{abs·surface·density}}+\text{abs·surface·density}\,\text{activator·conc}\right)-\frac{3.03\times10^{10}}{\text{activator·conc}+73\,\text{il2·conc}}$$

$$15169298.00-\frac{3.11\times10^{8}}{\text{activator·conc}}-\frac{\left(3.76\times10^{8}\right)\left(\sqrt{\text{abs·surface·density}}-\text{il2·conc}\right)}{\text{activator·conc}}+1.49\,(2\,\text{abs·surface·density}-\text{activator·conc}+\text{il2·conc})^{2}$$

$$-32552105.00+\frac{2.90\times10^{10}}{\text{abs·surface·density}}+30883.22\,\text{abs·surface·density}-3.38\,\text{abs·surface·density}\,\text{activator·conc}-\frac{48861030.00\left(4+\sqrt{\text{abs·surface·density}}\right)}{\text{activator·conc}\sqrt{\frac{\text{il2·conc}}{\text{activator·conc}}}}$$

$$15751220.00-222.07\,\text{abs·surface·density}+1.59\,(-3+2\,\text{abs·surface·density}-\text{activator·conc})^{2}-\frac{\left(4.02\times10^{8}\right)\left(\sqrt{\text{abs·surface·density}}-\text{il2·conc}\right)}{\text{activator·conc}}-781.22\,\text{il2·conc}^{2}$$

B.

$$1402298.40+\left(4.46\times10^{-4}\right)\text{abs·surface·density}^{3}+1626.15\,\text{activator·conc}-\frac{21061.46\,\text{abs·surface·density}}{\text{il2·conc}}$$

$$-1230176.60+\left(5.52\times10^{-4}\right)\text{abs·surface·density}^{3}+118975.62\sqrt{\text{activator·conc}}-\frac{11.41\,\text{abs·surface·density}^{2}}{\text{il2·conc}}$$

$$-1543881.00+\left(5.94\times10^{-4}\right)\text{abs·surface·density}^{3}+118188.48\sqrt{\text{activator·conc}}-\frac{\left(5.78\times10^{-3}\right)\text{abs·surface·density}^{3}}{\text{il2·conc}}$$

$$2395574.00+0.96\,\text{abs·surface·density}^{2}-\frac{6.82\times10^{8}}{\text{activator·conc}}+1074.62\,\text{activator·conc}-\frac{20141.44\,\text{abs·surface·density}}{\text{il2·conc}}$$

$$1573259.40+\left(4.80\times10^{-4}\right)\text{abs·surface·density}^{3}+1901.55\,\text{activator·conc}-\frac{24603.42\,\text{abs·surface·density}}{\text{il2·conc}}-13.88\,\text{activator·conc}\,\text{il2·conc}$$

$$-970660.23+\left(5.05\times10^{-4}\right)\text{abs·surface·density}^{3}+119624.65\sqrt{\text{activator·conc}}+\frac{10828456.00}{\text{il2·conc}}-\frac{27157.48\,\text{abs·surface·density}}{\text{il2·conc}}$$

$$2461761.70+\left(5.50\times10^{-4}\right)\text{abs·surface·density}^{3}-\frac{7.90\times10^{8}}{\text{activator·conc}}+1008.03\,\text{activator·conc}-\frac{11.31\,\text{abs·surface·density}^{2}}{\text{il2·conc}}$$

$$2521959.10+\left(5.17\times10^{-4}\right)\text{abs·surface·density}^{3}-\frac{476310.12\,\text{abs·surface·density}}{\text{activator·conc}}+1097.34\,\text{activator·conc}-\frac{20344.59\,\text{abs·surface·density}}{\text{il2·conc}}$$

$$-1056891.30+\left(5.25\times10^{-4}\right)\text{abs·surface·density}^{3}+118779.99\sqrt{\text{activator·conc}}-\frac{3267266.20}{\text{il2·conc}}-\frac{10.42\,\text{abs·surface·density}^{2}}{\text{il2·conc}}$$

$$-12665636.00+\left(2.07\times10^{-7}\right)\text{abs·surface·density}^{4}+6824096.50\,\text{activator·conc}^{1/8}-\frac{20028.20\,\text{abs·surface·density}}{\text{il2·conc}}$$

$$-995957.62+\left(5.14\times10^{-4}\right)\text{abs·surface·density}^{3}+118973.41\sqrt{\text{activator·conc}}-\frac{7613.08\,\text{abs·surface·density}}{\text{il2·conc}}-\frac{7.30\,\text{abs·surface·density}^{2}}{\text{il2·conc}}$$

$$4598624.60-5343.41\,\text{abs·surface·density}-\frac{574794.30\,\text{abs·surface·density}}{\text{activator·conc}}+1013.66\,\text{activator·conc}+1.28\,\text{abs·surface·density}^{2}\,\text{il2·conc}^{1/4}$$

$$1637049.10+\left(5.86\times10^{-4}\right)\text{abs·surface·density}^{3}+1156.41\,\text{activator·conc}-\frac{11.04\,\text{abs·surface·density}^{2}}{\text{il2·conc}}-\frac{\left(1.91\times10^{8}\right)\text{abs·surface·density}}{(\text{activator·conc}+\text{il2·conc})^{2}}$$

Figure S10: Individual symbolic regression equations in each ensemble for a) memory and b) CD4+ T cell yield responses.
NOTE: il2·conc = IL2 Concentration (U/ml), activator·conc = Carrier Concentration (carriers/ml), abs·surface·density = Antibody Surface Density (mAbs/μm$^2$)
